# Analysis of Essential Genes in *Clostridioides difficile* by CRISPRi and Tn-seq

**DOI:** 10.1101/2025.06.04.657922

**Authors:** Maia E. Alberts, Micaila P. Kurtz, Ute Müh, Jonathon P. Bernardi, Kevin W. Bollinger, Horia A. Dobrila, Leonard Duncan, Hannah M. Laster, Andres J. Orea, Anthony G. Pannullo, Juan G. Rivera-Rosado, Facundo V. Torres, Craig D. Ellermeier, David S. Weiss

## Abstract

Essential genes are interesting in their own right and as potential antibiotic targets. To date, only one report has identified essential genes on a genome-wide scale in *Clostridioides difficile*, a problematic pathogen for which treatment options are limited. That foundational study used large-scale transposon mutagenesis to identify 404 protein-encoding genes as likely to be essential for vegetative growth of the epidemic strain R20291. Here, we revisit the essential genes of strain R20291 using a combination of CRISPR interference (CRISPRi) and transposon-sequencing (Tn-seq). First, we targeted 181 of the 404 putatively essential genes with CRISPRi. We confirmed essentiality for >90% of the targeted genes and observed morphological defects for >80% of them. Second, we conducted a new Tn-seq analysis, which identified 346 genes as essential, of which 283 are in common with the previous report and might be considered a provisional essential gene set that minimizes false positives. We compare the list of essential genes to those of other bacteria, especially *Bacillus subtilis*, highlighting some noteworthy differences. Finally, we used fusions to red fluorescent protein (RFP) to identify 18 putative new cell division proteins, three of which are conserved in Bacillota but of largely unknown function. Collectively, our findings provide new tools and insights that advance our understanding of *C. difficile*.

**IMPORTANCE:** *Clostridioides difficile* is an opportunistic pathogen for which better antibiotics are sorely needed. Most antibiotics target pathways that are essential for viability. Here we use saturation transposon mutagenesis and gene silencing with CRISPR interference to identify and characterize genes required for growth on laboratory media. Comparison to the model organism *B. subtilis* reveals many similarities and a few striking differences that warrant further study and may include opportunities for developing antibiotics that kill *C. difficile* without decimating the healthy microbiota needed to keep *C. difficile* in check.

## INTRODUCTION

*Clostridioides difficile* infections (CDI) kill close to 13,000 people a year in the United States (1). Treating CDI is challenging because the antibiotics effective against *C. difficile* also impact the normal intestinal microbiota needed to keep *C. difficile* in check (2–4). There is a need for improved antibiotics that inhibit *C. difficile* more selectively. Most clinically useful antibiotics target proteins or pathways that are essential for viability, so a deeper understanding of the essential genes in *C. difficile* might provide foundational knowledge to guide antibiotic development. Essential genes are also interesting in their own right, as they provide insights into the most fundamental aspects of bacterial physiology.

Transposon sequencing (Tn-seq) identifies essential genes on a genome-wide scale based on the absence of insertions following saturation transposon mutagenesis (5, 6). However, several caveats must be kept in mind when interpreting the output of a Tn-seq experiment. For instance, insertion mutants that are viable but grow slowly will be lost from the mutant pool during outgrowth, so some apparently essential genes can be deleted. This caveat underscores the fact that binary categorization of genes as essential or nonessential is useful but an oversimplification. Tn-seq might also erroneously classify non-essential genes as essential due to polarity onto *bona fide* essential genes or because the random nature of Tn insertions means genes might be missed for stochastic reasons. Finally, Tn-seq does not provide insight into the actual function of essential genes because the phenotypic defects of the corresponding insertion mutants are not observed. Despite these caveats and limitations, Tn-seq is a powerful tool for prioritizing genes to investigate by more laborious methods.

CRISPR interference (CRISPRi) is a complementary approach for genome-wide interrogation of essential genes in bacteria (7–12). CRISPRi uses a single guide RNA (sgRNA) to direct a catalytically inactive Cas9 protein (dCas9) to a gene of interest, thereby repressing transcription (13). As the organism continues to grow and divide it becomes depleted of the targeted protein, potentially revealing phenotypic changes that precede cell death. Thus, CRISPRi provides functional information that Tn-seq cannot. However, CRISPRi shares with Tn-seq the problem of polarity, which has to be taken into consideration when interpreting phenotypes.

In 2015 Dembek et al. used Tn-seq to identify 404 protein-encoding genes as essential for vegetative growth in *C. difficile* strain R20291 on BHI media (14). As expected, most of these genes encode proteins involved in core biological processes and cell surface biogenesis, but some are of unknown function or not expected to be essential. Here, we revisit the essential genes of strain R20291 using a combination of CRISPRi and Tn-seq. First, we targeted 181 of the 404 putatively essential genes with CRISPRi to vet essentiality and identify terminal phenotypes. We confirmed essentiality for >90% of the targeted genes and observed morphological defects for >80% of them. Second, we conducted a new and more thorough Tn-seq analysis to identify genes essential for vegetative growth on TY media. We classified 346 protein-coding genes as essential, of which 283 (∼80%) were also essential in the previous study. Finally, we conducted a microscopy-based screen to identify potential cell division proteins. We discuss our findings in light of what is known about essential genes and cell division in other bacteria, particularly *Bacillus subtilis*.

## RESULTS AND DISCUSSION

### A library for CRISPRi knockdown of 181 putative essential genes

Our *C. difficile* CRISPRi plasmid has been described (15). It expresses *dCas9* from a xylose-inducible promoter (P_xyl_) and a sgRNA from a constitutively-active glutamate dehydrogenase promoter (P_gdh_). Constructing a knockdown library involved several steps: selecting the genes to be targeted, designing the sgRNAs, cloning those sgRNAs into the CRISPRi plasmid, and moving the finished plasmids from *E. coli* into *C. difficile* by conjugation. Because conjugation efficiencies are low, plasmids have to be moved from *E. coli* into *C. difficile* one-by-one. This step imposes a bottleneck that made it impractical to target all 404 essential genes identified previously. We therefore trimmed the gene list by excluding all transposon and phage-related genes (because these are not part of the core genome), most genes for tRNA synthetases and ribosomal proteins (to limit redundancy), and most genes for small proteins, defined here as fewer than 80 amino acids (240 nucleotides). Short genes are small targets for Tn insertion, so a disproportionate fraction are likely to be false positives. At this point we were left with 252 genes. Because CRISPRi is polar (7, 13, 16, 17), there is little to be gained by targeting multiple genes in an operon, so in most cases we targeted only one gene per transcription unit as annotated in BioCyc (v28.5, release Dec 2025) (18).

In the end, we selected a total of 181 putatively essential genes for CRISPRi knockdown (Table S1). We constructed a library of individual sgRNA clones, using two sgRNAs per gene for a total of 362 CRISPRi plasmids (Table S2). As negative controls, we constructed 20 CRISPRi plasmids with scrambled sgRNAs that do not target anywhere in the R20291 genome (Table S2). Plasmids were confirmed by sequencing across the P_gdh_::sgRNA element in *E.coli* and after conjugation into *C. difficile*. Of the genes targeted for knockdown, 86 have an essential ortholog in *Bacillus subtilis*, 62 have a non-essential ortholog in *B. subtilis*, and 33 have no *B. subtilis* ortholog, including four hypothetical genes. However, the number of genes of unknown function is larger than four because many of the non-hypotheticals have homology to domains with such broadly or ill-defined functions that it is not obvious what these genes do or why they would be essential (e.g., “glycosyltransferase,” “two-component response regulator,” or “DUF1846”).

Considering that 110 of the targeted genes are predicted to be in operons with other apparently essential genes, our study encompasses 281 of the 404 genes identified as essential by Tn mutagenesis, close to 70% of the total (14).

### Essentiality determined by CRISPRi knock-down largely agrees with Tn-seq data

The entire CRISPRi library was screened for viability defects by conducting spot titer assays on TY plates containing thiamphenicol at 10 μg/ml (hereafter TY-Thi10) and 1% xylose. Control plates lacked xylose. Using a 10-fold viability defect or small colony phenotype with at least one sgRNA as the cut-off, 167 of the 181 genes (92%) were confirmed as essential by CRISPRi, while 14 were not essential (Fig. 1A; Table S1). Similar results were obtained with both sgRNAs for 174 of the 181 genes tested (Table S1). None of the 20 non-targeting control sgRNAs caused a growth defect, indicating off-target effects are rare. We conclude that the vast majority of the genes Dembek et al. identified as essential by Tn-seq are also essential by CRISPRi (14).

**Fig. 1.**
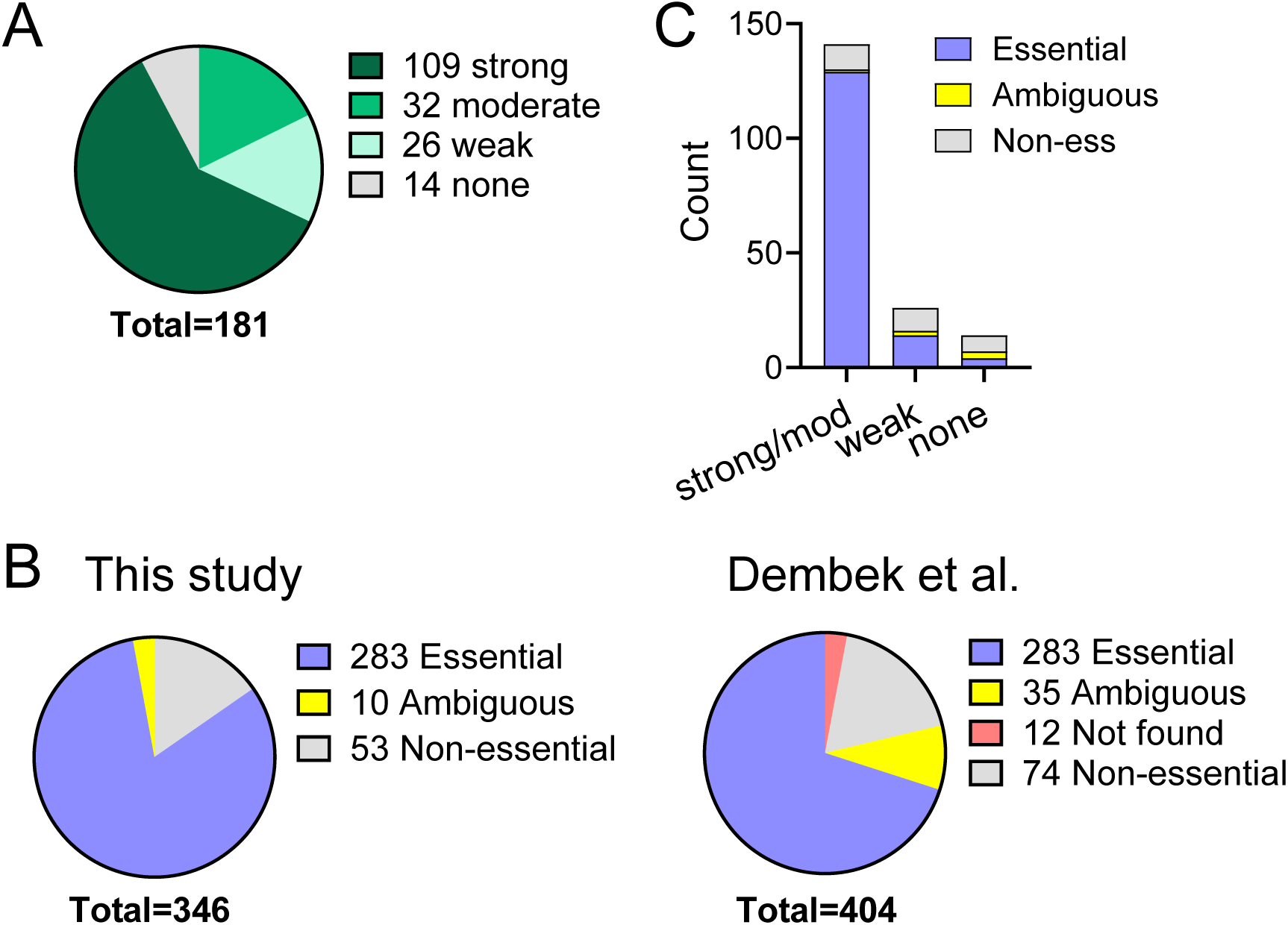
Summary of gene essentiality determined by CRISPRi and Tn-seq. (A) CRISPRi-induced viability defects were determined from spot titer assays on TY-thiamphenicol with 1% xylose. Viability defects were scored as strong (≥ 1000-fold), moderate (≥100-fold), weak (≥10-fold or full viability but colonies were small), or none (full viability, normal colony size). In cases where the two sgRNAs produced different results, the stronger viability defect was used. (B) Comparison of Tn-seq datasets. Of the 346 genes determined to be essential in our study, 283 were essential in the Dembek et al. set, and 10 were called ambiguous. Conversely, of the 404 Dembek et al. essential genes, 283 were essential in our dataset, 74 were non-essential, 12 were not found owing to use of different genome annotations, and 35 were ambiguous (Here, ambiguous combines three categories from Supplemental Table 3: unclear (11 genes), short (6 genes), unclear/non-essential (18 genes)). (C) Viability defects in CRISPRi correlate with likelihood a gene will be scored as essential by Tn-seq. Viability defects are from Table S1. Tn-seq calls come from Table S3.

### Terminal phenotypes due to CRISPRi knockdown of genes of known function

To look for morphological abnormalities that might facilitate provisional assignment of essential genes to functional pathways, cells were scraped from the last culture dilution that grew on the 1% xylose plates and examined by phase-contrast microscopy. As the project progressed, we added staining with FM4-64 to visualize the cytoplasmic membrane and Hoechst 33342 to visualize DNA. The morphological defects associated with CRISPRi silencing of all 181 genes are listed in Table S1.

CRISPRi knockdown of genes of known function often provoked expected morphological defects, such as filamentation in the case of cell division genes and aberrant nucleoid staining in the case of DNA replication genes (Fig. 2, Fig. S1, Table 1, Table S1). Also as expected, knockdown of DNA replication genes sometimes resulted in filamentation, presumably due to induction of the SOS response (19, 20). However, we also observed morphological defects that were not expected and are difficult to rationalize. For instance, knockdown of *rpoB* (β subunit of RNA polymerase) or *era* (GTPase involved in ribosome assembly) caused severe filamentation, while knockdown of *guaA* (synthesis of guanosine ribonucleotides) caused a mild chaining phenotype. To address whether the unexpected morphological abnormalities are an artifact of working with cells scraped from plates, we reexamined the filamentation phenotype of four non-division genes in broth about six doublings after inducing CRISPRi: *dnaH*, *rpoB*, *prfB* and *tilS*. We observed elongated cells in each case (Fig. S2). Thus, at least for this phenotype and these four genes, morphologies determined using plates are reliable.

**Fig. 2.**
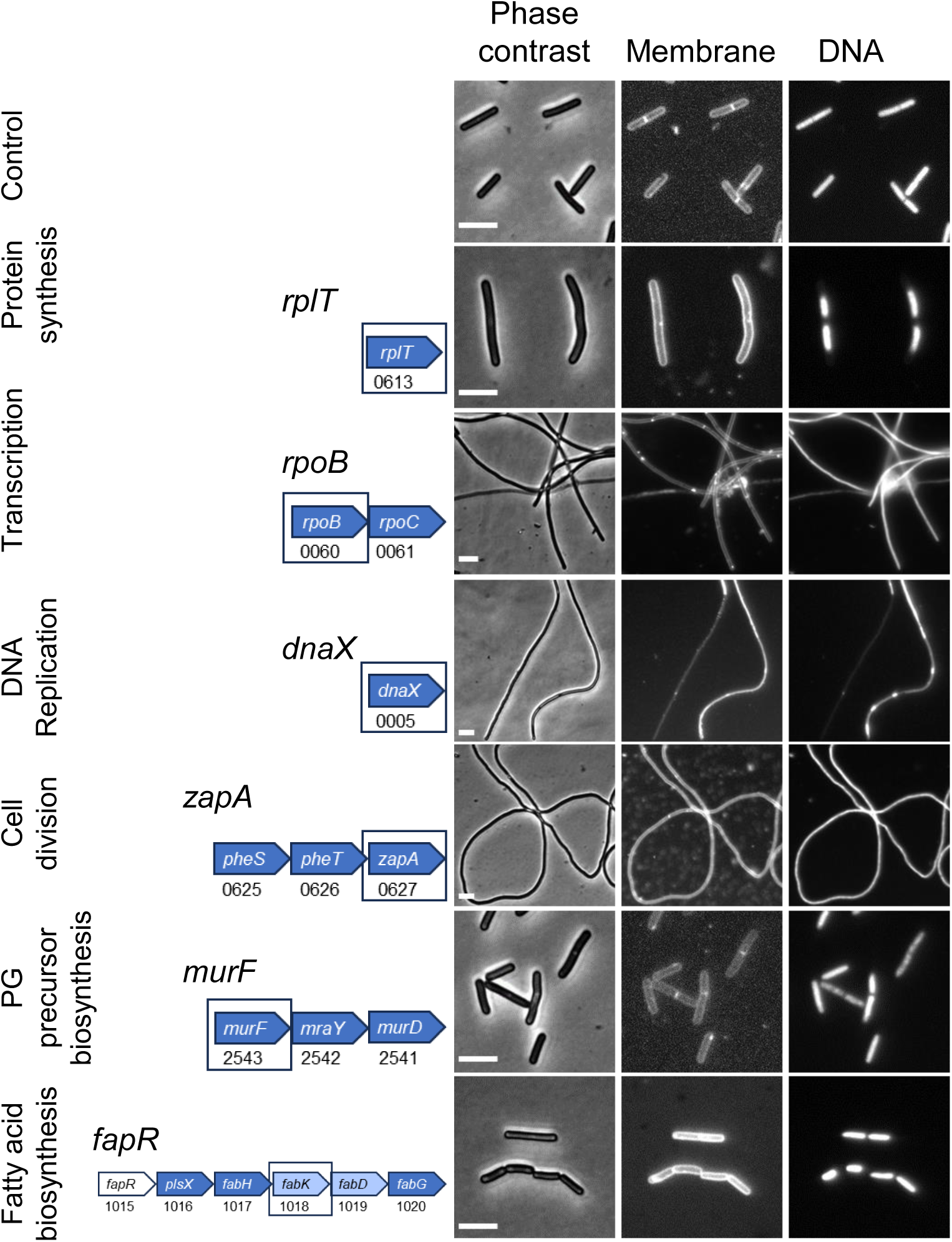
Morphology of CRISPRi strains with sgRNAs targeting genes in select functional pathways. Left: pathway. Middle: Predicted transcription unit. Targeted genes are boxed and indicated above the operon diagrams. Numbers are R20291 locus tags. Genes are color coded to indicate essentiality based on Tn-seq calls in Table S3. Blue: essential. Light blue: ambiguous. White: non-essential. Operon structure is not to scale. Right: Morphological changes based on phase contrast and fluorescence micrographs of cells scraped from viability plates. Membranes were stained with FM4-64 and DNA was stained with Hoechst 33342. Size bars are 5 µm. The control strain expressed an sgRNA that does not target anywhere in the genome. Micrographs are representative of at least two experiments. Figure S1 shows microscopy of more genes.

**Table 1.**
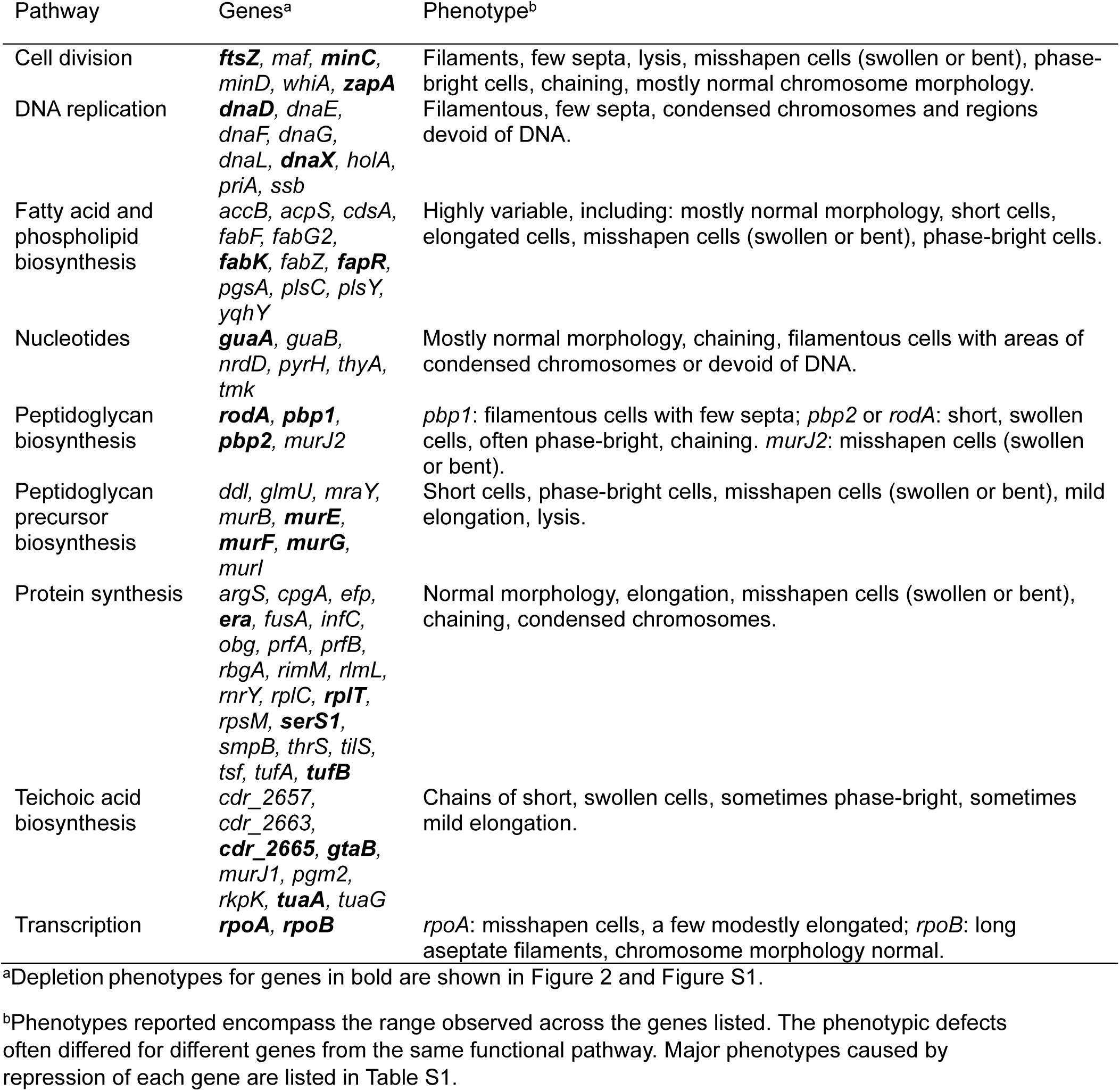
CRISPRi phenotypes of functional pathways.

Because morphological defects were only loosely associated with the function of well-studied genes, we conclude that CRISPRi is not sufficient for assigning genes of unknown function to physiological pathways. We are not the first to report unanticipated complexity among terminal phenotypes in a CRISPRi screen. For example, CRISPRi knockdown of the RNA polymerase gene *rpoC* and the phospholipid synthesis genes *psd* and *plsB* caused filamentation in *E. coli* (8). In addition, knockdown of multiple genes with no direct role in envelope biogenesis caused morphological defects in *B. subtilis* (7). These reports contrast with the narrower spectrum of morphological defects induced by antibiotics that target specific pathways (21–23). Antibiotics might be less subject to secondary effects because cells are visualized at early times after exposure and polarity is not an issue.

### Terminal phenotypes due to CRISPRi knockdown of genes of unknown function

Our CRISPRi library targeted 11 genes that could not be assigned to a functional category and were confirmed as essential in our own Tn-seq analysis as will be described below. CRISPRi caused a viability defect in nine cases, often accompanied by abnormal morphologies (Table 2). Examples include *cdr20291_0481* and *cdr20291_0828* (elongation), the *cdr20291_1053-1057* cluster (short, swollen, phase-bright cells and chaining), *cdr20291_1124* (chaining and many misshapen phase-bright cells) and *cdr20291_2526* (a few misshapen cells). The phenotype resulting from knockdown of *cdr20291_1124* could be due to reverse polarity onto the upstream gene *alaS*, which encodes an alanyl-tRNA synthetase. These genes warrant further investigation.

**Table 2.**
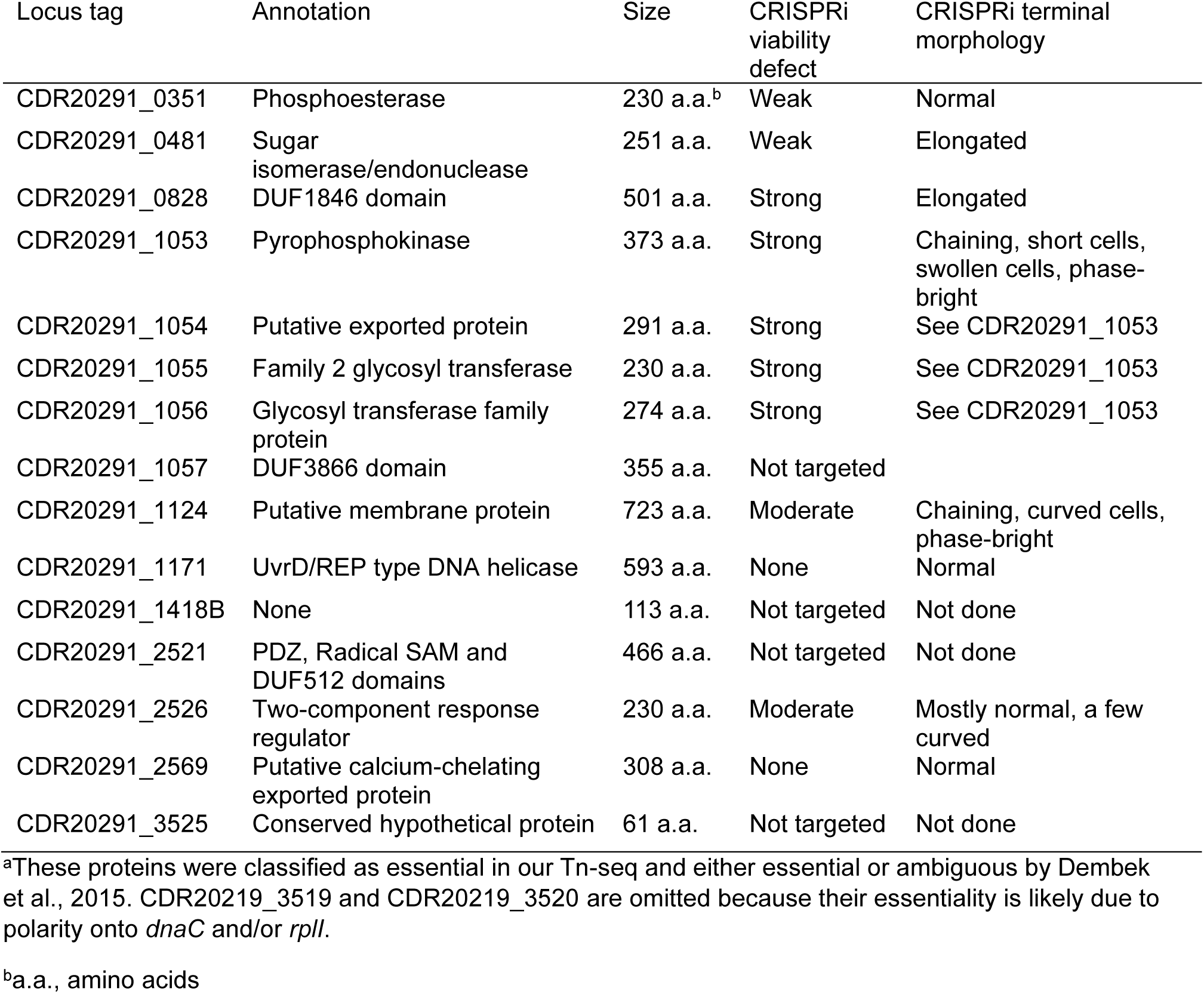
Essential genes not assigned to a physiological pathway^a^.

| Locus tag | Annotation | Size | CRISPRi viability defect | CRISPRi terminal morphology |
| --- | --- | --- | --- | --- |
| CDR20291_0351 | Phosphoesterase | 230 a.a.<sup>b</sup> | Weak | Normal |
| CDR20291_0481 | Sugar isomerase/endonuclease | 251 a.a. | Weak | Elongated |
| CDR20291_0828 | DUF1846 domain | 501 a.a. | Strong | Elongated |
| CDR20291_1053 | Pyrophosphokinase | 373 a.a. | Strong | Chaining, short cells, swollen cells, phase-bright |
| CDR20291_1054 | Putative exported protein | 291 a.a. | Strong | See CDR20291_1053 |
| CDR20291_1055 | Family 2 glycosyl transferase | 230 a.a. | Strong | See CDR20291_1053 |
| CDR20291_1056 | Glycosyl transferase family protein | 274 a.a. | Strong | See CDR20291_1053 |
| CDR20291_1057 | DUF3866 domain | 355 a.a. | Not targeted |  |
| CDR20291_1124 | Putative membrane protein | 723 a.a. | Moderate | Chaining, curved cells, phase-bright |
| CDR20291_1171 | UvrD/REP type DNA helicase | 593 a.a. | None | Normal |
| CDR20291_1418B | None | 113 a.a. | Not targeted | Not done |
| CDR20291_2521 | PDZ, Radical SAM and DUF512 domains | 466 a.a. | Not targeted | Not done |
| CDR20291_2526 | Two-component response regulator | 230 a.a. | Moderate | Mostly normal, a few curved |
| CDR20291_2569 | Putative calcium-chelating exported protein | 308 a.a. | None | Normal |
| CDR20291_3525 | Conserved hypothetical protein | 61 a.a. | Not targeted | Not done |

<sup>a</sup>These proteins were classified as essential in our Tn-seq and either essential or ambiguous by Dembek et al., 2015. CDR20219\_3519 and CDR20219\_3520 are omitted because their essentiality is likely due to polarity onto *dnaC* and/or *rplI*.
<sup>b</sup>a.a., amino acids

### Rationale for decision to conduct Tn-seq

As noted above, our CRISPRi screen was based on the only published genome-wide analysis of gene essentiality in *C. difficile*. That study made essentiality calls based on a single pool of mutants containing ∼77,000 unique Tn insertions plated on BHI media (14). We reasoned that an independently-derived list of essential genes based on multiple biological replicates and larger insertion libraries would serve as a useful resource to the *C. difficile* community. We also thought a new Tn-seq dataset might serve as a “tie-breaker” for the 14 putatively essential genes that did not appear to be essential by CRISPRi, i.e., failure to recover insertions in those genes would suggest our sgRNAs were ineffective, while recovery of insertions would suggest the genes are non-essential and were missed in the previous study for stochastic reasons.

### Generation of Tn insertion libraries and identification of essential genes

We used the same R20291 strain and mariner-based transposon as in the previous study (14). Mariner is a good choice for *C. difficile* because it inserts at TA dinucleotides and the genome G+C content is 29% (24, 25). However, our experimental design differed from Dembek et al. in three noteworthy respects: (i) we used TY media, (ii) we constructed three independent insertion libraries, (iii) and we determined insertion profiles at both an early and a late timepoint because gradual loss of slow-growing mutants from the pools influences perceptions of gene essentiality. Our early timepoint consisted of primary insertion libraries recovered directly from selection plates after ∼18 hours of incubation. For a later timepoint, libraries were sub-cultured in duplicate into TY and harvested after seven generations of outgrowth.

Insertion profiles were analyzed using TRANSIT2 and the *C. difficile* R20291 reference genome NC_013316.1 (26, 27). Depending on the experimental replicate, insertions were identified in 117,217 to 204,061 of the 502,945 unique TA dinucleotides in the R20291 genome (Table 3). A total of 289,505 TA sites sustained at least one Tn insertion across the three libraries. TRANSIT2 makes essentiality calls by comparing the observed frequency of Tn insertions to the availability of potential TA insertion sites. Genes are classified as essential (E or EB, depending on the model for statistical analysis), not essential (NE), or unclear (U) (28). Genes with too few TA sites for statistical analysis are designated S. After inspecting the output from TRANSIT2, we manually reclassified eleven NE or U genes as essential, giving them the designation Ei for “essential by inspection.” Ten of these genes had a large number of TA sites but very few insertions. An example is the tRNA-synthetase *valS* (CDR20291_3114), with insertions in only four of the possible 266 TA dinucleotides after outgrowth (Table S3A). For comparison, TRANSIT2 scored the cell division gene *ftsZ* as essential even though there were insertions in three out of 110 TA sites. All ten genes that we moved to Ei based on few insertions are considered essential in *C. difficile* and *B. subtilis* (14, 29). The final Ei gene, *murJ2* (CDR20291_3335), had a large number of insertions but almost all of these were at the 3’ end of the gene (Fig. 3A). *murJ2* was previously classified as essential in *C. difficile* by Dembek et al. but its ortholog is not essential in *B. subtilis* due to functional redundancy (29, 30).

**Fig. 3.**
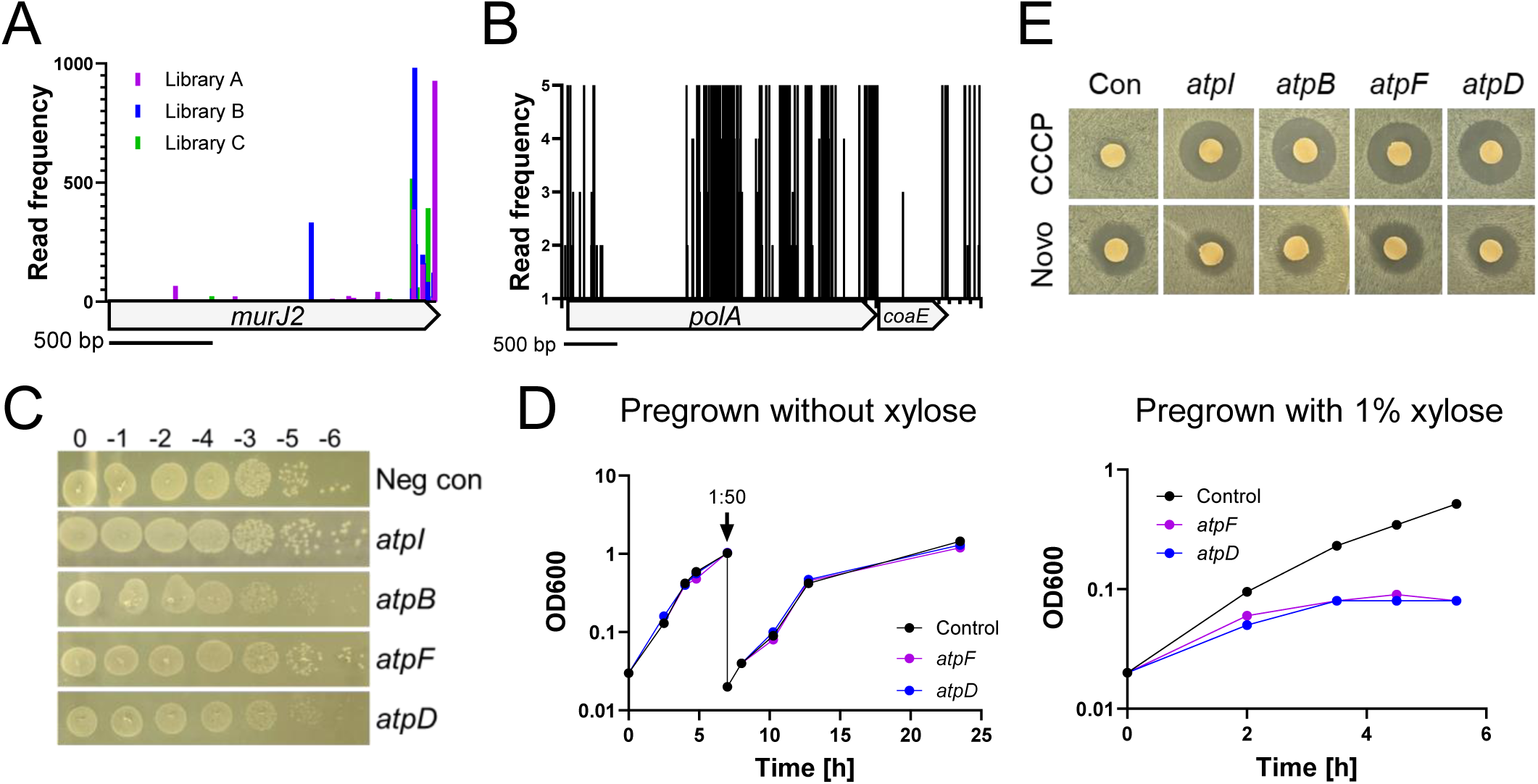
Essentiality follow-up. (A) Transposon insertion profile for *murJ2*. Vertical lines represent mapped insertion sites and are scaled to indicate the number of sequence reads mapping to that site. Although *murJ2* sustained numerous insertions, ∼80% were in the last 10% of the gene, suggesting the non-essentiality call by TRANSIT2 is incorrect. (B) Transposon insertion profile for *polA* indicating that only the N-terminal domain is essential. Read frequency was scaled to 5 to highlight the absence of reads in the N-terminal domain. The average number of reads per *polA* site with at least one read was 173. (C) Spot titer assays of CRISPRi strains targeting genes in the *atp* operon. Serial dilutions of overnight cultures were spotted on TY-Thi10 plates with 1% xylose. Plates were imaged after incubation at 37°C for ∼18h. Silencing *atpB* and *atpD* resulted in small colonies, while growth after silencing *atpI* and *atpF* was comparable to the negative control. (D) Pre-depletion of ATP synthase proteins impairs growth. Starter cultures were grown overnight in TY-Thi10 without (left) or with (right) 1% xylose, then subcultured into TY-Thi10 with 1% xylose and growth was followed by measuring optical density at 600 nm. To prolong growth, cultures in the left panel were back-diluted at 7h. (E) Zone of inhibition assays reveal CRISPRi knockdown of the *atp* operon increases sensitivity to CCCP. Plates were imaged after incubation at 37°C for ∼18h. Novobiocin (Novo) served as a control. Guides in panels C-E were: *atpI* (5531), *atpB* (5583), *atpF* (5581), *atpD* (5579) or a negative control that does not target anywhere in the gemone.

**Table 3.** Number of unique transposon insertions from experimental replicates^a^.

| Replicate | Unique TA sites hit | Fraction of total TA sites |
| --- | --- | --- |
| Library A | 178,325 | 0.355 |
| Library B | 168,519 | 0.335 |
| Library C | 143,213 | 0.285 |
| Combined Libraries A-C | 289,505 | 0.576 |
| Outgrowth A1 | 135,217 | 0.269 |
| Outgrowth A2 | 117,217 | 0.233 |
| Outgrowth B1 | 127,947 | 0.254 |
| Outgrowth B2 | 135,056 | 0.269 |
| Outgrowth C1 | 167,894 | 0.334 |
| Outgrowth C2 | 204,061 | 0.406 |

<sup>a</sup>Total TA sites = 502,945

Of the 3673 annotated protein-coding genes in R20291, 346 were scored as essential for vegetative growth in the initial libraries and/or after outgrowth (Table S3A). We grouped these genes into functional categories similar to those used in previous studies of *B. subtilis* and *S. aureus* (Table 4; Table S3B) (31, 32). As expected, over half are involved in DNA metabolism (25 genes), RNA metabolism (24 genes), protein synthesis (113 genes) or cell envelope biogenesis (76 genes). Also as expected, the majority of *C. difficile’s* essential genes are conserved; BioCyc assigned a *B. subtilis* ortholog for 272 of the 346 genes, of which 169 are essential (Table S3A, B) (29).

**Table 4.**
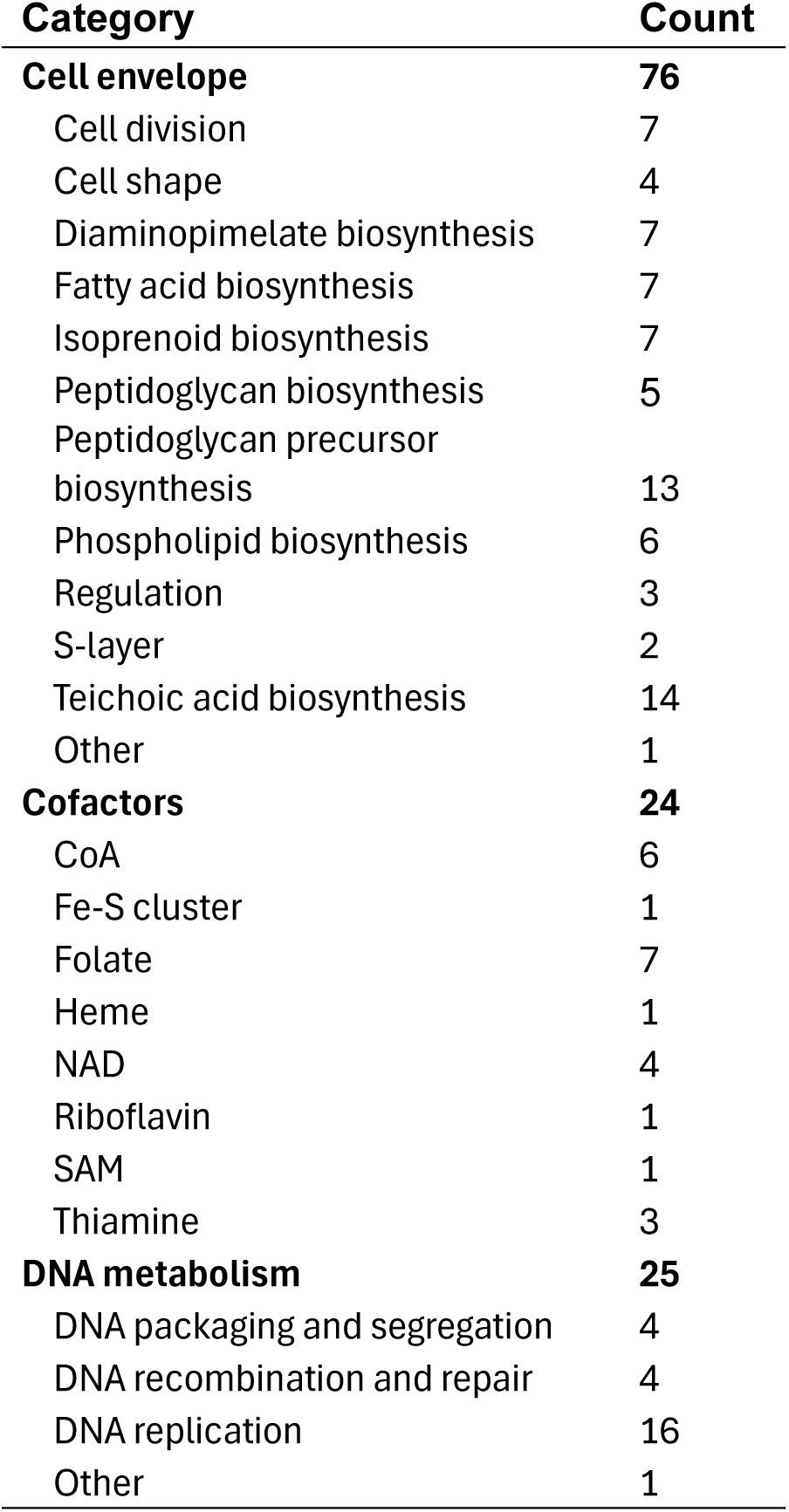
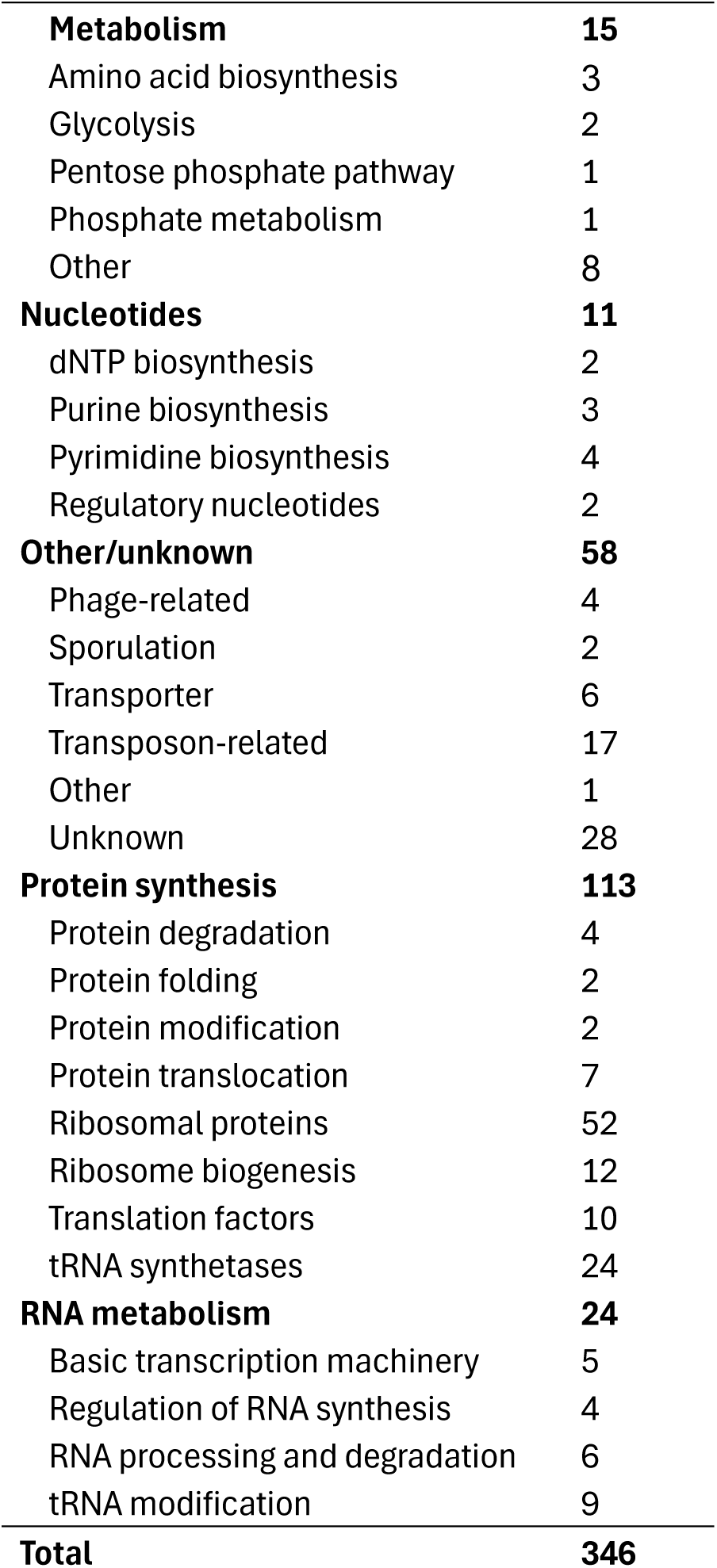
Tn-seq essential genes by category.

| **Category** | **Count** | **Category** | **Count** |
| --- | --- | --- | --- |
| **Cell envelope** | **76** | **Metabolism** | **15** |
| Cell division | 7 | Amino acid biosynthesis | 3 |
| Cell shape | 4 | Glycolysis | 2 |
| Diaminopimelate biosynthesis | 7 | Pentose phosphate pathway | 1 |
| Fatty acid biosynthesis | 7 | Phosphate metabolism | 1 |
| Isoprenoid biosynthesis | 7 | Other | 8 |
| Peptidoglycan biosynthesis | 5 | **Nucleotides** | **11** |
| Peptidoglycan precursor biosynthesis | 13 | dNTP biosynthesis | 2 |
| Phospholipid biosynthesis | 6 | Purine biosynthesis | 3 |
| Regulation | 3 | Pyrimidine biosynthesis | 4 |
| S-layer | 2 | Regulatory nucleotides | 2 |
| Teichoic acid biosynthesis | 14 | **Other/unknown** | **58** |
| Other | 1 | Phage-related | 4 |
| **Cofactors** | **24** | Sporulation | 2 |
| CoA | 6 | Transporter | 6 |
| Fe-S cluster | 1 | Transposon-related | 17 |
| Folate | 7 | Other | 1 |
| Heme | 1 | Unknown | 28 |
| NAD | 4 | **Protein synthesis** | **113** |
| Riboflavin | 1 | Protein degradation | 4 |
| SAM | 1 | Protein folding | 2 |
| Thiamine | 3 | Protein modification | 2 |
| **DNA metabolism** | **25** | Protein translocation | 7 |
| DNA packaging and segregation | 4 | Ribosomal proteins | 52 |
| DNA recombination and repair | 4 | Ribosome biogenesis | 12 |
| DNA replication | 16 | Translation factors | 10 |
| Other | 1 | tRNA synthetases | 24 |
|  |  | **RNA metabolism** | **24** |
|  |  | Basic transcription machinery | 5 |
|  |  | Regulation of RNA synthesis | 4 |
|  |  | RNA processing and degradation | 6 |
|  |  | tRNA modification | 9 |
|  |  | **Total** | **346** |

### Comparison of our Tn-seq data to Dembek et al. and to CRISPRi

There is good overall agreement between our Tn-seq essentiality calls and those made previously. Of the 346 genes identified as essential in our experiments, 283 (82%) were also essential for Dembek et al. (Fig. 1B; Table S3A). As expected based on the larger size of our insertion libraries, we scored fewer genes as essential, 346 versus 404 (Fig. 1B). Our list of essential genes includes 53 considered nonessential by Dembek et al., while those investigators identified 121 essential genes that did not make our cutoffs. Of these, 12 encode proteins that are not annotated in the genome sequence we used and were thus invisible to our analysis. An additional 6 were scored as S and 29 as U or U/NE. That leaves 74 *bona fide* discrepancies, genes that were essential for Dembek et al. but not essential for us. Several of these differences will be discussed below, but the most likely explanations have to do with statistical cut-offs that factor into essentiality calls and the stochastic nature of Tn mutagenesis.

There is also good overall agreement between our CRISPRi and Tn-seq data sets. Of the 141 genes for which CRISPRi elicited a strong or moderate viability defect, 129 (∼90%) scored as essential in our Tn-seq (Fig. 1C; Table S3A). Conversely, only 4 out of 14 genes (∼30%) that appeared to be nonessential by CRISPRi nevertheless scored as essential in our Tn-seq. These four genes are an uncharacterized DNA helicase (CDR20291_1171), a sporulation-associated phosphatase (*ptpB*), an acetyl-CoA thiolase (*thlA2*), and a putative exported Ca^2+^-chelating protein (*ykwD*). None of these has an essential ortholog in *B. subtilis*. Two labs have constructed null mutants of *ptpB*, indicating it is not essential (33, 34). One study reported a growth defect (34), which might explain why *ptpB* appears to be essential by Tn-seq.

### DNA metabolism

Some DNA replication proteins have different names in *B. subtilis* and *E. coli*. Where there are conflicts, we adopted the names used in *B. subtilis*, which in some cases differ from the names used in BioCyc. We identified 16 widely conserved DNA replication genes as essential in *C. difficile*. All but *pcrA* and *polA* were previously classified as essential in *C. difficile*, and all but *polA* is essential in *B. subtilis* (14, 29). Interestingly, *polA* is domain essential in *C. difficile*—Tn insertions were recovered in the C-terminal 3’ to 5’ exonuclease and DNA polymerase domains but not in the N-terminal 5’ to 3’ exonuclease domain, which removes Okazaki fragments (Fig. 3B). Similar restricted essentiality of the *polA* 5’ to 3’ exonuclease domain has been reported in *Streptococcus* and *Haemophilus* (35, 36). Organisms like *B. subtilis* in which the entire *polA* gene is dispensable have an RNAse H that can remove Okazaki fragments (37).

Interestingly, *C. difficile* lacks *dnaB* (38). DnaB is an essential protein in *B. subtilis*, where it works together with DnaD and DnaI to load the replicative helicase DnaC onto *oriC* DNA (39). DnaB and DnaD are structurally related. It has been proposed that in *C. difficile* the DnaD ortholog (CDR20291_3512) fulfills the functions of both DnaB and DnaD (38).

*C. difficile* has four essential DNA packaging and segregation genes, all of which are also essential in *B. subtilis*. In addition, there are three essential DNA recombination and repair genes, none of which are essential in *B. subtilis*.

LexA, which represses genes involved in the SOS response, is required for viability in *C. difficile* but not in *B. subtilis*. A *C. difficile lexA* Clostron insertion mutant has been described and grows poorly, so its apparent essentiality by Tn-seq may be due to slow growth rather than lack of viability *per se* (19). However, the strong viability defect we observed upon CRISPRi knockdown of *lexA* (Table S1) raises the possibility that the reported mutant retains partial function or acquired a suppressor.

### RNA metabolism

As expected, the core subunits and major sigma factor (σ^70^) of RNA polymerase are all essential. Surprisingly, the omega subunit (*rpoZ*) is also essential according to Tn-seq, even though it is not essential in *B. subtilis*, *S. aureus* or *E. coli* (29, 40, 41). The apparent essentiality of *rpoZ* is likely to be an artifact of polarity because it is predicted to be co-transcribed with three widely conserved essential genes: *dapF*, *gmk*, *coaBC*. The elongation factor *greA* and three termination/anti-termination factors (*nusA*, *nusG* and *rho*) are essential. Of these, only *nusA* is essential in *B. subtilis* (29, 42). In *C. difficile rho* mutations have been reported, including an early frameshift, but the gene could not be deleted, possibly because the mutant is too sick (43).

Fifteen genes for enzymes that modify RNA were essential in our analysis, of which twelve were essential or ambiguous for Dembek et al., but only eight are considered essential in *B. subtilis*. Most of these genes encode proteins needed to generate mature tRNAs or rRNAs from precursor transcripts.

### Protein synthesis

There are 55 annotated ribosomal proteins in BioCyc (Dec 18, 2024), 52 of which scored as essential by our Tn-seq. Most of these were also identified as essential by Dembek et al. and are essential in *B. subtilis*. Instances of ribosomal protein genes scored as essential in *C. difficile* but as non-essential in *B. subtilis* could reflect polarity. Five widely conserved small GTPases involved in ribosome assembly are essential, as are ten translation factors, including *smpB*, which encodes a component of the SsrA tagging complex that rescues stalled ribosomes by trans-translation (44). We confirmed essentiality of *smpB* by CRISPRi (Table S1). The essentiality of *smpB* is unlikely to be an artifact of polarity because it is not predicted to be co-transcribed with any other genes. SmpB is essential in *S. aureus* (32, 45) but not in *E. coli*, *Streptococcus sanguinis*, or *B. subtilis* (29, 46, 47). An interesting omission from the list of essential translation factors is elongation factor Tu (EF-Tu), which is essential in *B. subtilis* (29). This difference can be explained by the presence of two EF-Tu genes in *C. difficile*, *tufA* and *tufB*, which are 100% identical at the DNA level. Simultaneous knockdown of *tufA* and *tufB* with CRISPRi caused a strong viability defect, demonstrating EF-Tu is indeed required for viability (Table S1).

We identified 24 essential tRNA synthetases, all of which are also essential according to Dembek et al. There are several noteworthy differences in comparison to *B. subtilis*. First, synthetases for asparagine (*asnS*), threonine (*thrS*) and tyrosine (*tyrS*) are essential in *C. difficile* but not *B. subtilis*, which has alternative routes for generating the corresponding charged tRNAs (48–50). Second, although *glnS* is essential in *C. difficile*, this gene does not exist in *B. subtilis* or most other gram-positive bacteria, which generate Gln-tRNA^Gln^ by a different route. Namely, *C. difficile* charges tRNA^Gln^ directly with glutamine, as in *E. coli*, while most Gram-positive bacteria generate glutaminyl-tRNA^Gln^ by (mis)charging tRNA^Gln^ with glutamate, which is then amidated to glutamine (51, 52). Lastly, *C. difficile* has two annotated genes for ligating proline to tRNA^pro^, the essential gene *proS1* (CDR20291_0038) and the non-essential gene *proS2* (CDR20291_0039). According to RNA-sequencing, both are expressed during vegetative growth (53). *B. subtilis* has only a single *proS* gene, which is essential and more similar to *C. difficile proS1* than *proS2*.

Five proteases appear to be important for viability in *C. difficile*: *clpX*, *htrA*, *lon, prp* and the M16 family protease *cdr20291_1161*. Of these, only *prp* is essential in *B. subtilis*. Prp is a cysteine protease needed to remove an N-terminal extension from ribosomal protein L27 (54).

The apparent essentiality of *lon* and *cdr20291_1161* in *C. difficile* are likely to be artifacts of polarity onto *engB* and *dapG*, respectively. ClpX is a component of the ClpXP protease complex, one of the major housekeeping proteases in bacteria (55). *C. difficile* has only one *clpX* gene but two genes for ClpP, which might explain why *clpX* is essential but *clpP1* and *clpP2* are not. HtrA proteases are involved in protein quality control (56). TRANSIT2 scored *htrA* as essential despite a high number of Tn insertions (67 out of 127 TA sites) and this gene was not essential for Dembek et al. In bacteria, protein synthesis begins with *N*-formyl methionine (fMet). Peptide deformylase (*def*) and methionine aminopeptidase (*map*) are essential enzymes that work sequentially to remove the formyl group from about 90% of proteins and the initiating methionine from about half of proteins. *E. coli* has only one *def* and one *map* gene, both of which are essential (57). *C. difficile* has two predicted *map* genes and two predicted *def* genes. Of these, only *map1* is essential by Tn-seq. This situation is reminiscent of *B. subtilis*, which also has two *def* and two *map* genes. The *def* genes are functionally redundant and at least one must be present for viability (58, 59). The essentiality of the *map* genes *in B. subtilis* is less clear. One study found *mapA* is essential but *mapB* is not (60), while another found neither is individually essential (29).

Bacteria have a plethora of systems for exporting proteins out of the cytoplasm, of which the three most important are the General Secretion (Sec) system, the Twin Arginine Translocation (Tat) system, and the Signal Recognition Particle (SRP) system (61). There is no Tat system in *C. difficile*, but the genes for the Sec and SRP systems are present and essential. The Sec system uses an ATPase named SecA to power export of proteins through a membrane channel composed of SecEYG. Interestingly, *C. difficile* has two *secA* paralogs, which handle different protein substrates and are both essential (62). The SRP system works together with SecEYG to integrate proteins into the cytoplasmic membrane. Three genes associated with the SRP system (*ffh*, *ftsY* and *srpM*) were scored as essential, although the apparent essentiality of *srpM* might result from polarity onto *ffh*; *srpM* is not essential in *B. subtilis*.

### Cell envelope

Numerous genes involved in membrane biogenesis are essential in *C. difficile*. An unexpected exception is the *accBCDA* gene cluster for synthesis of malonyl-CoA, the substrate for fatty acid synthesis. This result is difficult to explain and probably incorrect because the *acc* cluster is essential according to Dembek et al. and we confirmed essentiality by CRISPRi (Table S1). Moreover, *acc* genes are also essential in *B. subtilis* (29). Nevertheless, the *acc* cluster sustained numerous Tn insertions in our study (e.g., ten of the 48 TA sites in *accB*, the first gene in the operon). We identified three membrane biogenesis genes that are essential in *C. difficile* but not in *B. subtilis*: *fabH*, *yqhY*, and *gpsA*. The *fabH* discrepancy can be explained by the presence of two *fabH* genes in *B. subtilis* (63). *B. subtilis* Δ*yqhY* mutants are not stable (64), implying *yqhY* is quasi essential in that organism. Regarding *gpsA*, although Koo et al. reported it is dispensable in *B. subtilis* (29), an earlier study found it is essential (65), which agrees with what we see in *C. difficile*.

*C. difficile* synthesizes isoprenoids via the methylerythritol (MEP) pathway (66). Accordingly, *dxr* and *ispDEFGH* were all essential by Tn-seq. Isoprenoids are essential in bacterial because they are precursors for quinones and carrier lipids such as undecaprenyl phosphate (Und-P) required for synthesis of peptidoglycan and teichoic acids (67). *C. difficile* lacks quinones (68) so the essentiality of the MEP pathway presumably reflects the importance of Und-P. Consistent with this inference, the predicted undecaprenyl pyrophosphate synthase UppS1 is essential, although that conclusion comes with a caveat because insertions in *uppS1* are probably polar onto the essential phospholipid biosynthesis gene *cdsA* (69). Interestingly, *C. difficile* has a non-essential *uppS* paralog called *uppS2* that might be involved in synthesis of the wall teichoic acid PS-II (70). UppS2 is not essential by Tn-seq, and RNA-sequencing implies expression of *uppS2* is ∼60-fold lower in vegetative cells compared to *uppS1* (53).

The *C. difficile* cell has a unique proteinaceous surface-layer (S-layer) and a unique wall teichoic acid, PS-II, whose structure is very different from the wall teichoic acids of other Gram-positive bacteria (71). Both the S-layer and PS-II are essential by Tn-seq, although the existence of (unhealthy) null mutants of *slpA* indicates the S-layer is not strictly required for viability (72, 73). Multiple studies point to essentiality of PS-II (70, 74, 75). Whether PS-II is essential because it plays a critical role in cell envelope integrity or because disruption of the PS-II gene cluster depletes the pool of Und-P needed for peptidoglycan synthesis remains to be determined (76, 77).

The universal precursor for peptidoglycan synthesis is lipid II, a disaccharide-pentapeptide attached to Und-P (78). As expected, many lipid II genes are essential, including six *dap* genes for biosynthesis of lysine and diaminopimelic acid, and nine *mur* genes for various steps in lipid II assembly. Lipid II is transported across the cytoplasmic membrane by flippases, of which there are two known families, MurJ and Amj (30, 79). BLAST searches indicate *C. difficile* lacks Amj but has two MurJ orthologs, both of which are essential. MurJ1 is part of the PS-II gene cluster and proposed to transport a lipid-linked precursor for PS-II synthesis (74), which leaves MurJ2 as the likely lipid II flippase for peptidoglycan synthesis. Some non-essential proteins distantly related to MurJ can be identified using HHPred and could also potentially transport lipid II (80, 81). Further work is needed to establish the functions of the two clear MurJ paralogs and rule out the presence of alternative or additional lipid II transporters (30, 82, 83).

The final steps of peptidoglycan synthesis involve incorporation of new disaccharide-pentapeptide subunits into the existing wall by sequential glycosyltransferase (GTase) and transpeptidase (TPase) reactions (84, 85). These reactions are catalyzed by two types of penicillin-binding proteins (PBPs) (86). Class A PBPs (aPBPs) are bifunctional enzymes with both a GTase domain and a TPase domain, while class B PBPs (bPBPs) have a TPase domain and form a complex with a SEDS-family GTase (87–89). *C. difficile* encodes one aPBP (PBP1), three bPBPs (PBP2, PBP3, and SpoVD), and two SEDS proteins (RodA and SpoVE). Of these proteins, we confirmed by Tn-seq that PBP1, PBP2 and RodA are essential for vegetative growth (14). Although *spoVE* was also classified as essential, it sustained Tn insertions in about half the available TA sites (Table S3A) and the gene has been deleted previously (90). In confirmation and extension of previous reports (15, 91), CRISPRi knockdown of PBP1 caused filamentation, while CRISPRi knockdown of PBP2 and RodA resulted in formation of short, swollen, phase-bright cells, with some chaining (Fig. S1). These morphologies implicate PBP1 in cell division and PBP2 in elongation, respectively. We also examined red fluorescent protein (RFP) fusions to the PBPs and observed that both localize to division sites (Fig. 4). Septal localization of PBP1 has been reported by Shen’s group, who showed it is the primary synthase for septal peptidoglycan (91). Septal localization of PBP2 suggests the RodA/PBP2 complex might also contribute to cell division, as further suggested by the mild chaining phenotypes caused by CRISPRi knockdown. Both RFP-PBP1 and RFP-PBP2 exhibited some fluorescence along the cell cylinder, which could indicate they contribute to elongation, especially in the case of PBP2. However, localization to the cell cylinder is not diagnostic of a function in elongation because this is the default location of divisome proteins when they are not at the septum. Finally, it should be noted that non-canonical 3-3 crosslinks made by L,D-transpeptidases (LDTs) are essential for vegetative growth in *C. difficile*, but none of the five LDTs in the *C. difficile* genome is individually essential owing to functional redundancy (92).

**Fig. 4.**
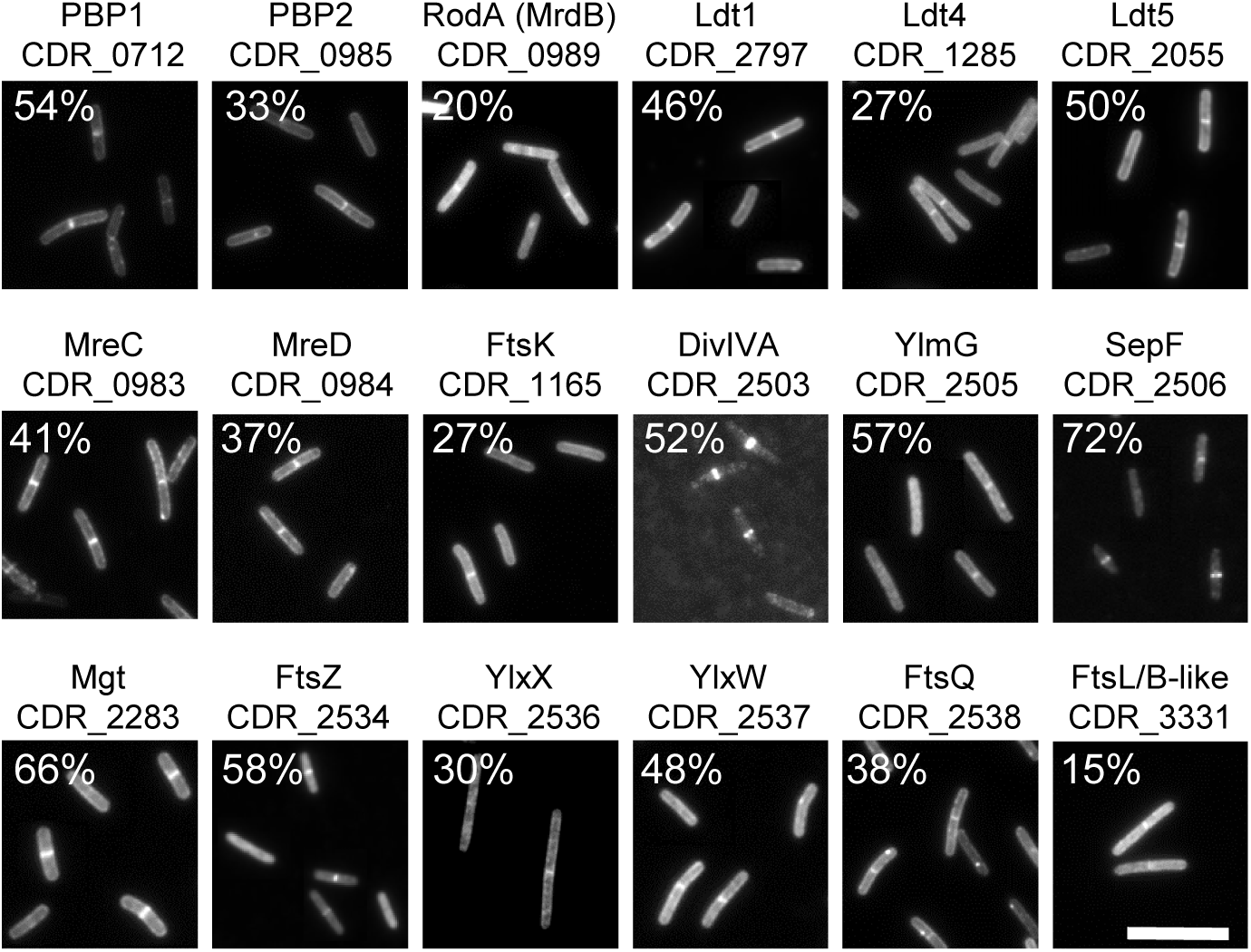
Representative fluorescence micrographs of fixed cells that produced the indicated proteins fused to RFP. Percentages indicate the fraction of cells scored positive for septal localization (n ≥ 202 cells). Space bar = 10 µm.

Our Tn-seq identified two cell envelope-related regulatory loci as essential: *walRK* and *ddlR*. These regulators were also essential for Dembek et al. *walRK* is a two-component system known to be essential for cell wall homeostasis and viability in numerous Bacillota, including *C. difficile* (53, 93). DdlR is essential for peptidoglycan synthesis because it activates expression of the D-alanyl-D-alanine ligase *ddl* (94).

### Cell shape and division

In rod-shaped bacteria, the essential peptidoglycan synthases work in the context of loosely defined complexes known as the elongasome and the divisome (84, 85). The *C. difficile* elongasome appears to comprise the RodA/PBP2 bipartite peptidoglycan synthase and four Mre proteins (MreB1, MreB2, MreC and MreD). All of these are essential by Tn-seq and CRISPRi, although this inference will need to be revisited with non-polar deletions. CRISPRi knockdown implicates these genes primarily in elongation, because the predominant terminal morphologies include short, swollen cells with some chaining (Fig. S1, Table S1).

Among canonical divisome proteins, only *ftsZ* and its assembly factors *sepF* and *zapA* are essential in *C. difficile*. Neither *sepF* nor *zapA* is essential in *B. subtilis* (95–97). The greater importance of *sepF* and *zapA* in *C. difficile* might be due to the absence of an *ftsA* ortholog (98). As noted above, the primary septal peptidoglycan synthase is the class A enzyme PBP1 (91). Consistent with that inference, CRISPRi against *pbp1* induces filamentation, however additional morphological defects such as bending and chaining suggest PBP1 might contribute to elongation as well (15). Curiously, the division site placement genes *minCDE* are essential in *C. difficile*. This result might be an artifact of polarity onto the essential SEDS gene *rodA*, because the Min system is not essential in *B. subtilis* or most other bacteria (99). Tn-seq identified *maf* as essential. Maf is a nucleotide pyrophosphatase whose overproduction causes filamentation in both *B. subtilis* and *E. coli*, but Maf is not essential in either organism (100–102). The DNA-binding protein WhiA was essential for Dembek et al. and we observed a weak viability defect and modest cell elongation by CRISPRi, but *whiA* is not essential in our Tn-seq experiments. WhiA is conserved in monoderms and essential in *Mycobacterium tuberculosis* but not *Streptomyces* or *B. subtilis*, where it has been linked to cell division and chromosome segregation (103–107).

### Use of RFP fusions to identify new divisome proteins

We have a long-standing interest in bacterial cell division, so we extended our studies to include a screen for divisome proteins (108–112). Using CRISPRi knockdown to identify divisome proteins by screening for a filamentous phenotype comes with two major caveats—polarity onto a *bona fide* division gene will generate false positives and depletion of non-essential divisome proteins might not cause cells to become longer than normal. A more direct approach is to use fluorescent tags to screen for proteins that localize to the division site. Here the major caveat is that the tag might interfere with proper localization. We used BLAST searches to identify homologs of known morphogenesis proteins, which were fused to a codon-optimized red fluorescent protein (RFP) and produced from a plasmid under control of the xylose-inducible promoter, P_xyl_ (15). Some of these proteins are encoded in (predicted) operons with proteins of unknown function, so we constructed RFP fusions to several of these as well. Although septal localization is strong evidence for a role in cell division, lack of septal localization is uninformative because we did not test whether our RFP fusions are functional. We screened a total of 25 proteins, of which 18 localized and are discussed below (Fig. 4). The seven that did not localize are MreB1, MreB2, FtsL, FtsB, SpoVE, CDR_3330, and CDR_2504.

Seven enzymes for peptidoglycan synthesis exhibited convincing midcell localization, including the two essential PBPs (PBP1 and PBP2), one essential SEDS protein (RodA), one non-essential monofunctional glycosyltransferase related to PBPs (Mgt), and three non-essential LDTs (Ldt1, Ldt4 and Ldt5). Of these, PBP1 was already known to localize to sites of cell division (91), but septal localization of the remaining enzymes is new and suggests they too contribute to synthesis of septal peptidoglycan. Somewhat surprisingly, the canonical elongasome proteins MreC and MreD localized strongly to the midcell, even though our fusions to MreB1 and MreB2 did not. Mre proteins have been reported to localize transiently at or near the midcell in a few other bacteria (113–116). Further work is warranted to investigate the role of the Mre proteins in *C. difficile* and the possibility that MreC and MreD localize independently of MreB, for which there is precedent from non-rod-shaped bacteria that have MreC and MreD but lack MreB (116, 117).

*C. difficile* orthologs of five widely-conserved divisome proteins localized to the midcell: FtsZ, FtsK, FtsQ, SepF, DivIVA, as did CDR_3331, a unique protein with limited structural similarity to both FtsL and FtsB, which in *C. difficile* are used for asymmetric division during sporulation (14, 91). Septal localization of *C. difficile* FtsZ has been reported previously (118). Septal localization of FtsQ is new but probably misleading because *C. difficile ftsQ* is a sporulation gene and not expressed during vegetative growth (14, 53, 90), whereas we produced RFP-FtsQ from P_xyl_. Immediately downstream of *ftsQ* are two genes of unknown function, *ylxW* and *ylxX*, that according to RNA-sequencing are expressed in vegetative cells (53). YlxW and YlxX are encoded downstream of *ftsQ* in many Bacillota and have been proposed on this basis to play a role in envelope biogenesis (119). Our observation that these proteins localize to the midcell argues they are involved in cell division. Another novel divisome protein identified in our screen is YlmG, a small membrane protein encoded in the *sepF* operon of many Gram-positive bacteria and Cyanobacteria (98). Mutants of *ylmG* have been constructed in several organisms and exhibit thin septa, poor sporulation, and/or aberrant nucleoid compaction and segregation, depending on the species (98). In closing, and for completeness, we note that four additional proteins have been shown previously to localize to the division site in *C. difficile*: ZapA, MldA, MldB and MldC (112, 120). This brings total number of documented divisome proteins to 22.

### Metabolism

For an insightful overview of energy metabolism in *C. difficile*, readers are referred to a review by Neumann-Schaal et al. (121). Briefly, *C. difficile* is an obligate anaerobe that generates energy through fermentation of sugars and amino acids, the latter by a process known as Stickland reactions (122, 123). There is no electron transport chain. Hence, the five genes that are essential for menaquinone biosynthesis in *B. subtilis* are not found in *C. difficile’s* genome. The TCA cycle is incomplete and is used to generate precursor metabolites rather than energy. Fermentation pathways generate ATP directly by substrate level phosphorylation but can also be used via electron bifurcation and the Rnf complex to generate a motive force across the cytoplasmic membrane (124, 125). Whether this is a proton or a sodium-ion motive force is not yet known; we will assume protons for simplicity, i.e., a PMF*. C. difficile* has an F_0_F_1_-type ATP synthase, which, depending on the needs of the organism, can consume the PMF to generate ATP or hydrolyze ATP to generate a PMF.

Few of the genes involved in these various pathways scored as essential by Tn-seq. Genes for the TCA cycle, acetate kinase, and the major Stickland reductases for glycine, proline and leucine are all non-essential, as are the genes for the RNF complex and three electron bifurcation complexes (*etf* genes). The essentiality of genes for glycolysis is less clear because eight of these were essential for Dembek et al. but only two (*eno*, *tpiA*) were essential in our experiments. Glycolysis might have more of a contribution on BHI, which contains glucose, than on TY. Differences in slow growth and statistical cutoffs that impact essentiality calls may also factor into the discrepancies. In support of this explanation, we observed a small colony phenotype when we used CRISPRi to knock down expression of four glycolysis genes (*fba*, *gapB*, *pgi*, and *pfkA*) that were essential for Dembek et al. but not in our Tn-seq (Table S1). A further point to keep in mind is that glycolysis genes could be more important for supplying precursor metabolites rather than energy in *C. difficile*.

A noteworthy discrepancy concerns the ten gene operon for the F-type ATPase. Dembek et al. scored nine of the genes as essential, but all ten were non-essential in our Tn-seq experiments. This gene cluster is too large to have escaped Tn insertions by chance. The most likely explanation for this discrepancy has to do with how slow growth affects perceptions of essentiality because we observed that CRISPRi knockdown of *atpB* and *atpD* resulted in a small colony phenotype (Fig. 3C). We also tested the effect of knockdowns in TY broth using one sgRNA that caused a small colony phenotype (*atpD*) and one that did not (*atpF*). Interestingly, both knockdowns caused a strong growth defect, but only if cultures were pre-grown overnight in 1% xylose to deplete the AtpD or AtpF proteins before sub-culturing (Fig. 3D). As an aside, we found that all four *atp* operon knockdowns were sensitized to subinhibitory concentrations of the uncoupler carbonyl cyanide m-chlorophenylhydrazone (CCCP, Fig. 3E), which hints at the potential for using our CRISPRi library to study drug targets in *C. difficile* (7, 12).

Three genes (*hisC*, *ilvB* and *ilvC*) involved in amino acid biosynthesis were identified as essential despite utilization of growth medium rich in tryptone. These genes were not essential for Dembek et al. Note, however, that there are seven essential lysine biosynthesis genes, which we categorized under cell envelope rather than metabolism owing to their role in synthesis of diaminopimelate for peptidoglycan. Finally, the global regulator CodY is essential by Tn-seq. CodY is widely conserved in Bacillota and senses GTP and branched chain amino acids to regulate gene expression in response to the energetic and nutritional needs of the cell. In *C. difficile* CodY represses hundreds of genes during exponential growth, and a *codY* null mutant grows poorly upon entry into stationary phase (126–128), which likely explains the Tn-seq result.

### Nucleotides and cofactors

We identified eleven genes essential for nucleotide biosynthesis. All eleven were also essential or ambiguous for Dembek et al. and five are essential in *B. subtilis* as well. One interesting difference is that an anaerobic ribonucleotide reductase encoded by *nrdD* and *nrdG* is essential in *C. difficile*, but these genes are not found in *B. subtilis*, which has instead an aerobic ribonucleotide reductase encoded by *nrdE* and *nrdF* that are not found in *C. difficile* (129). Two additional exceptions are *guaA* (GMP synthase) and *thyA* (thymidylate synthase), which are essential in *C. difficile* and *S. aureus* but not *B. subtilis* (14, 29, 32). Two genes for regulatory nucleotides appear to be essential in *C. difficile*, the cyclic-di-GMP phosphodiesterase *yybT* and the bifunctional (pp)pGpp synthase/hydrolase *relA*. Essentiality of *relA* was confirmed by CRISPRi (Table S1). The *B. subtilis* paralogs of these genes are not essential (29). Essentiality of *yybT* is likely to be an artifact of polarity onto *rplI* or *dnaC*, but the genes transcribed with *relA* are not essential. In *C. difficile relA* is called *rsh* and synthesizes exclusively pGpp (130, 131). As an aside, we note that cyclic-di-AMP is essential in *C. difficile* growing on rich media, but c-di-AMP synthases were not identified by Tn-seq because there are two of them, neither of which is individually essential (132).

Twenty-four genes are essential for synthesis of cofactors despite utilization of media containing tryptone and yeast extract. All but two of these were also essential or ambiguous for Dembek et al., and 14 have an essential ortholog in *B. subtilis*. Curiously, neither we nor Dembek et al. scored dihydrofolate reductase (*dfrA*) as essential. Dihydrofolate reductase is the target of several important antibiotics and essential in *E. coli*, *B. subtilis*, *S. sanguinis*, and *S. aureus* (29, 32, 46, 47).

### Phage and Transposon-related genes

The *C. difficile* genome has a remarkably high content of mobile genetic elements (25, 133). Mobile genetic elements are not part of the core genome and thus should not be essential for viability. Nevertheless, twenty-one genes classified as essential appear to reside on a prophage or a transposon. Some of these might be false positives because only eight were also essential or ambiguous for Dembek et al. Even the eight genes classified as essential in both studies are likely due to indirect effects such as induction of a lytic prophage.

### Transporters

Six genes for transporters were classified as essential in our Tn-seq, three of which were also essential for Dembek et al. and were confirmed by CRISPRi (Table S1). These encode a predicted Ktr potassium transporter and a predicted CorA-like divalent metal ion transporter. In *B. subtilis*, there are two Ktr systems, which are not essential but improve growth at high osmolarity (134).

### Sporulation

Curiously, both we and Dembek et al. classified the sporulation-associated phosphatases *ptpA* and *ptpB* as ambiguous or essential for vegetative growth. Two labs have reported null mutants of these genes, so they are not formally essential (33, 135, 136). Loss of *ptpA* or *ptpB* enhances sporulation, which we confirmed using CRISPRi against *ptpB* (Table S1). We presume that *ptp* genes are essential by Tn-seq because enhanced sporulation reduces vegetative growth.

### Genes of unknown function

Our Tn-seq analysis identified 28 putatively essential genes that could not be assigned to a functional pathway. None of these genes have an essential ortholog in *B. subtilis*, although in five cases BioCyc identified a non-essential ortholog. Eleven of these genes were not essential for Dembek et al. and in two cases (*cdr20291_3519* and *cdr20291_3520*) essentiality is likely due to polarity onto *rplI or dnaC*. That leaves 15 genes that are essential or ambiguous in two independent Tn-seq studies and are therefore likely to be *bona fide* essential genes. As noted above in the discussion of our CRISPRi experiments, we silenced expression of eleven of these genes and observed a viability defect for nine of them, often accompanied by abnormal morphologies (Table 2, Table S1). The apparently essential genes of unknown function constitute a high value gene set from the perspectives of bacterial physiology and antibiotic development.

### Conclusions

In summary, we identified 346 protein-encoding genes that by Tn mutagenesis are essential for vegetative growth of *C. difficile* strain R20291 on TY media. Of these, 283 were also identified as essential by Tn mutagenesis in a previous study (14) and 169 have an essential ortholog in *B. subtilis* (29). Overall, these results are broadly consistent with studies of gene essentiality in model organisms such as *E. coli*, *B. subtilis* and *S. aureus* (29, 32, 45, 46, 57). The 283 *C. difficile* genes identified as essential in two independent Tn mutagenesis studies can be regarded as a consensus “essentialome” that minimizes false positives. Most of these genes play key roles in foundational cellular processes such as DNA replication, transcription, translation and cell envelope biogenesis. But the consensus essentialome also includes 15 genes that could not be assigned to any functional pathway (Table 2, Table S3A, B). These genes might be targets for antibiotics that kill *C. difficile* without decimating the healthy microbiota needed to keep *C. difficile* in check.

We also used CRISPRi knockdown to investigate 181 genes that had been identified as essential in a previous Tn-seq analysis (14). Our goals were to vet essentiality and screen for morphological defects that would facilitate assigning genes of unknown function to physiological pathways. Our CRISPRi platform used a plasmid that expresses *dCas9* from a xylose-inducible promoter (P_xyl_) and an sgRNA from a strong constitutive promoter (P_gdh_) (15). CRISPRi resulted in reduced plating efficiencies and/or small colony phenotypes on TY-xylose plates for 167 of the 181 genes targeted, a very high confirmation rate of 92%. The 14 genes for which no viability defect was observed could be false positives from the previous report or genes for which our sgRNAs were ineffective. Of these genes, ten sustained insertions in our Tn-seq experiments, so we infer they are non-essential. Four did not sustain Tn insertions and are therefore likely to be essential genes that were poorly repressed by our sgRNAs. Importantly, no growth defects were observed using 20 control sgRNAs that did not target anywhere in the genome, indicating off-target effects are rare.

Microscopy of surviving cells scraped from the TY-xylose plates revealed most knockdowns resulted in morphological abnormalities (151 out of 181 genes, 83%). Disappointingly, however, the utility of these defects for making functional assignments was limited by the observation that repressing genes of known function often resulted in non-intuitive defects. For example, repressing RNA polymerase gene *rpoB* resulted in severe filamentation suggestive of a cell division defect, while repressing the nucleotide biosynthesis gene *guaA* caused a chaining phenotype suggestive of a daughter cell separation defect. Non-intuitive phenotypes have also been reported in other CRISPRi screens (7, 8).

The findings and resources presented here should help guide future studies of *C. difficile*. First, our results can be used to prioritize genes for more rigorous but labor-intensive investigation using depletion strains with in-frame deletions (137). The 15 apparently essential genes that could not be assigned to a functional pathway seem like a good place to start. Second, our CRISPRi library can be leveraged to investigate antibiotic sensitivities (7, 12, 138), which might illuminate gene function and reveal vulnerabilities that can be exploited to improve treatment of *C. difficile* infections. Third, the identification of 18 proteins that localize to the midcell raises new questions related to *C. difficile* morphogenesis. For example, septal localization of the canonical elongation proteins MreC and MreD suggests they contribute to cell division and/or *C. difficile* elongates by inserting new peptidoglycan near the midcell. In addition, our discovery that YlmG, YlxW and YlxX localize to the division site provides the most direct evidence to date that these conserved but enigmatic proteins play a role in cell division.

## METHODS

### Strains, media, and growth conditions

Most bacterial strains used in this study are listed in Table S4. Strains and plasmids constructed for the CRISPRi library are summarized separately in Table S2. *C. difficile* strains were derived from R20291 (139). *C. difficile* was routinely grown in tryptone-yeast extract (TY) medium, supplemented as needed with thiamphenicol at 10 μg/ml (TY-Thi10). TY medium consisted of 3% tryptone, 2% yeast extract, and 2% agar (for plates). Brain heart infusion (BHI) media was prepared per manufacturer’s (DIFCO) instructions. *C. difficile* strains were maintained at 37°C in an anaerobic chamber (Coy Laboratory Products) in an atmosphere of 2% H_2_, 5% CO_2_, and 93% N_2_. *Escherichia coli* strains were grown in LB medium at 37°C with chloramphenicol at 10 μg/ml and/or ampicillin at 100 μg/ml as needed. LB medium contained 1% tryptone, 0.5% yeast extract, 0.5% NaCl, and 1.5% agar (for plates). OD_600_ measurements were made with the WPA Biowave CO8000 tube reader in the anaerobic chamber.

### Plasmid and strain construction

Plasmids are listed in Table S5 and were constructed with HiFi DNA Assembly from New England Biolabs (Ipswich, MA). Oligonucleotide primers (Table S6) were synthesized by Integrated DNA Technologies (Coralville, IA). CRISPRi plasmids were constructed as described in (15). Regions constructed by PCR were verified by DNA sequencing. Plasmids were propagated in *E. coli* HB101/pRK24 and conjugated into *C. difficile* R20291 according to (53). Final R20291 CRISPRi strains were verified by PCR amplifying and sequencing the guide region. Details relevant to other plasmid construction are provided in Table S5.

### CRISPRi screen

Overnight cultures grown in TY-Thi10 were serially diluted 10-fold in TY, and 5 µL spotted on TY-Thi10 and TY-Thi10 1% (w/v) xylose plates. Plates were incubated at 37°C overnight and imaged the following morning (∼18 h). Cells were scraped from select spots (usually the last spot with growth) and resuspended in 50 µL TY. Cell suspensions were supplemented with 5 µg/mL FM4-64 (red fluorescent membrane stain, Thermo Scientific) and 15 µg/mL Hoechst 33342 (blue fluorescent DNA stain, Invitrogen) and imaged by phase-contrast and fluorescence microscopy.

### Protein localization

R20291 harboring plasmids that expressed RFP-tagged proteins under xylose control were grown in TY-Thi10 overnight, subcultured into TY-Thi10 with 0.1% or 1% xylose, grown to an OD_600_ of about 0.6, and fixed with 4% buffered paraformaldehyde as described (53, 120, 140). Fixed cells were photographed under phase-contrast and (red) fluorescence. Septal localization was scored manually by inspecting cells for the presence of a fluorescent band near the midcell. MicrobeJ was used to keep track of cells scored positive or negative for septal localization (141).

### Microscopy

Cells were immobilized using thin agarose pads (1% w/v agarose). Phase-contrast micrographs were recorded on an Olympus BX60 microscope equipped with a 100× UPlanApo objective (numerical aperture, 1.35). Micrographs were captured with a Hamamatsu Orca Flash 4.0 V2+ complementary metal oxide semiconductor (CMOS) camera. Excitation light was generated with an X-Cite XYLIS LED light source. Red fluorescence was detected with the Chroma filter set 49008 (538 to 582 nm excitation filter, 587 nm dichroic mirror, and a 590 to 667 nm emission filter). Blue fluorescence was detected with the Olympus filter set U-MWU (330-385 nm excitation filter, 400 nm dichroic mirror and a 420 nm barrier emission filter).

### Transposon library construction

Plasmid pRPF215 is a quasi-suicide plasmid that harbors the *Himar1 mariner* transposase gene under control of P_tet_ (14). The gene for TetR does not have a terminator and transcription reads through into the origin of replication, presumably disrupting plasmid replication. Addition of anhydrotetracycline therefore both induces the transposase and causes plasmid loss. A single colony of R20291/pRPF215 was used to inoculate a 2 mL overnight culture in TY-Thi10. Twenty independent overnight cultures were grown for each transposon library construction. After overnight growth, each was then sub-cultured 1:50 into 2 mL TY and grown to an OD_600_ of 0.3. From each subculture an aliquot was removed and spread on to two large (15 cm diameter) plates of TY agar with 80 µg/mL lincomycin (RPI) and 100 ng/mL anhydrotetracycline (Sigma), for a total of 40 plates. We used higher concentrations of lincomycin than originally published (14) because we found 80 µg/mL lincomycin decreased the number of false positives. The amount of subculture to plate was experimentally determined to give roughly 5000-8000 colonies. Typically, we used 220 µL of subculture diluted with TY to 600 µL, a volume suitable for spreading evenly on a large plate. A dilution series of one subculture was also plated on TY to calculate plating efficiency. Selection plates typically grew one colony for every 500 plated (i.e., an efficiency of about 2 x 10^-3^). Plates were incubated for 20 hours at 37°C. Cells were then scraped off the plates with 5 mL TY each, pooled, amended to 10% DMSO, aliquoted and stored at –80°C. This material was referred to as the primary transposon library. Suspensions of the primary libraries typically had an OD_600_ of about 6. The concentration of viable cells was quantitated by plating aliquots on TY plates and was typically around 3 x 10^8^ CFU/mL. Three independent libraries were constructed on different days.

### Tn-seq sample preparation

DNA samples were prepared directly from 1 mL of primary library or from 10 mL culture that had been grown for an additional 7 doublings in TY. To avoid creating a bottleneck, 10 mL TY was inoculated with 2.2 x 10^7^ CFU. There are 502,945 possible TA insertion sites in the R20291 chromosome, thus cultures were started with a ratio of about 45 CFU per TA site. DNA libraries for Illumina sequencing were prepared based on modifications of Karash et al. (142). Briefly, regions adjacent to any transposon insertion were amplified by single primer extension. The resulting products were extended with a cytosine-tail, which then allowed further amplification by PCR. The upstream primer recognizes the transposon sequence, incorporates the P5 sequence for Illumina sequencing and a sample-specific barcode; the downstream primer recognizes the C-tail and incorporates the P7 sequence.

Genomic DNA was prepared using the Monarch Genomic DNA purification kit from NEB, using the protocol for Gram positive bacteria. A maximum of 2 x 10^9^ cells were pelleted. Lysis was facilitated through the addition of 0.5 mg hen egg white lysozyme (Boehringer Mannheim) and 20 U mutanolysin (Sigma), and DNA was eluted in 35 µL with a typical yield of 200 ng/µL. Linear extension PCR was performed on 100 ng DNA in 50 µL with Taq polymerase (NEB) and primer Tn-ermB-2 (anneal: 30 s at 55°C, extend 30 s at 68°C, 50 cycles). The resulting product was spin-column purified (Zymo Research Clean & Concentrator kit) and eluted in 12 µL. A C-tail was added by extending with terminal transferase (NEB) in a 20 µL reaction, using 1.25 mM dCTP (NEB) and 50 µM ddCTP (MilliporeSigma/Roche). The product was again spin-column purified and eluted in 10 µL. Final PCR amplification used 1 µL of C-tailed DNA in a 35 µL reaction mixture, Taq polymerase and primers P7-16G and P5-Tn-Px (x: variable barcode; anneal: 30 s at 62°C, extend 30 s at 68°C, 35 cycles). The resulting product was separated on a 1.5% agarose gel in Tris Acetate EDTA buffer (TAE). Fragments of 300-500 base pair length were excised, purified with the Zymo Research Gel DNA recovery kit, and eluted in 10 µL. DNA concentration was quantitated with the Qubit dsDNA assay and was typically around 5 ng/µL. Four samples with distinct barcodes were combined and submitted for sequencing (Illumina HiSeq X, 150-bp PE reads) with Admera Health Biopharma Services (South Plainfield, NJ). Samples were spiked with 5% PhiX DNA to improve data quality.

### Sequencing data processing

Raw sequencing files were first trimmed with Trimmomatic to eliminate poor quality reads (143). The first four bases before the barcodes were then removed using Trim Sequences and the resulting files were de-multiplexed using the Barcode splitter, both on Galaxy (144). Reads were aligned to the reference genome of R20291 (NC_013316.1 or ASM2710v1) using the Burrows-Wheeler Aligner (BWA) provided in TRANSIT (27). Finally, the resulting Wig files were compared in TRANSIT2 which evaluates gene essentiality both by Gumbel analysis and binomial analysis (145). The former makes essentiality calls based on insertion gaps, i.e. consecutive TA sites lacking transposon insertions, using the Gumbel distribution (146). The latter calls essentiality for small genes lacking insertions which can be difficult to detect by the more conservative Gumbel algorithm (28). Essentiality calls are either “E” when identified by Gumbel or “EB” when identified by the Binomial analysis. Table S3 lists genes that were called essential in primary insertion libraries using cells scraped from plates, or after an additional 7 generations of growth. The library dataset was generated from three independently constructed transposon libraries. The outgrowth dataset was generated from two independent growth cultures from each of the three independent libraries. We present both the separate data output as well as a combined essentiality call (Table S3). The latter was further hand-edited by including 11 genes (indicated as “Ei” for “essential by inspection”) that appeared to have mistakenly called non-essential by TRANSIT2. Ten of these genes had very few insertions despite numerous possible TA sites, while the eleventh had a large number of insertions but mostly at the 3’ end of the gene.

## Acknowledgements

This work was supported by Public Health Service Grants R21 AI159071 (D.S.W) and R01 AI155492 (C.D.E. and D.S.W). A.J.O. and F.V.T. were supported by NSF REU DBI-1852070. H.M.L. and J.G.R.-R. were supported by NSF REU DBI-2244169. We thank members of the Ellermeier and Weiss laboratories for helpful discussions, John Cronan for information on *gpsA* and Erin Purcell for information on *relA*.

