## Supplemental Figures for "Analysis of Essential Genes in *Clostridioides difficile* by CRISPRi and Tn-seq"

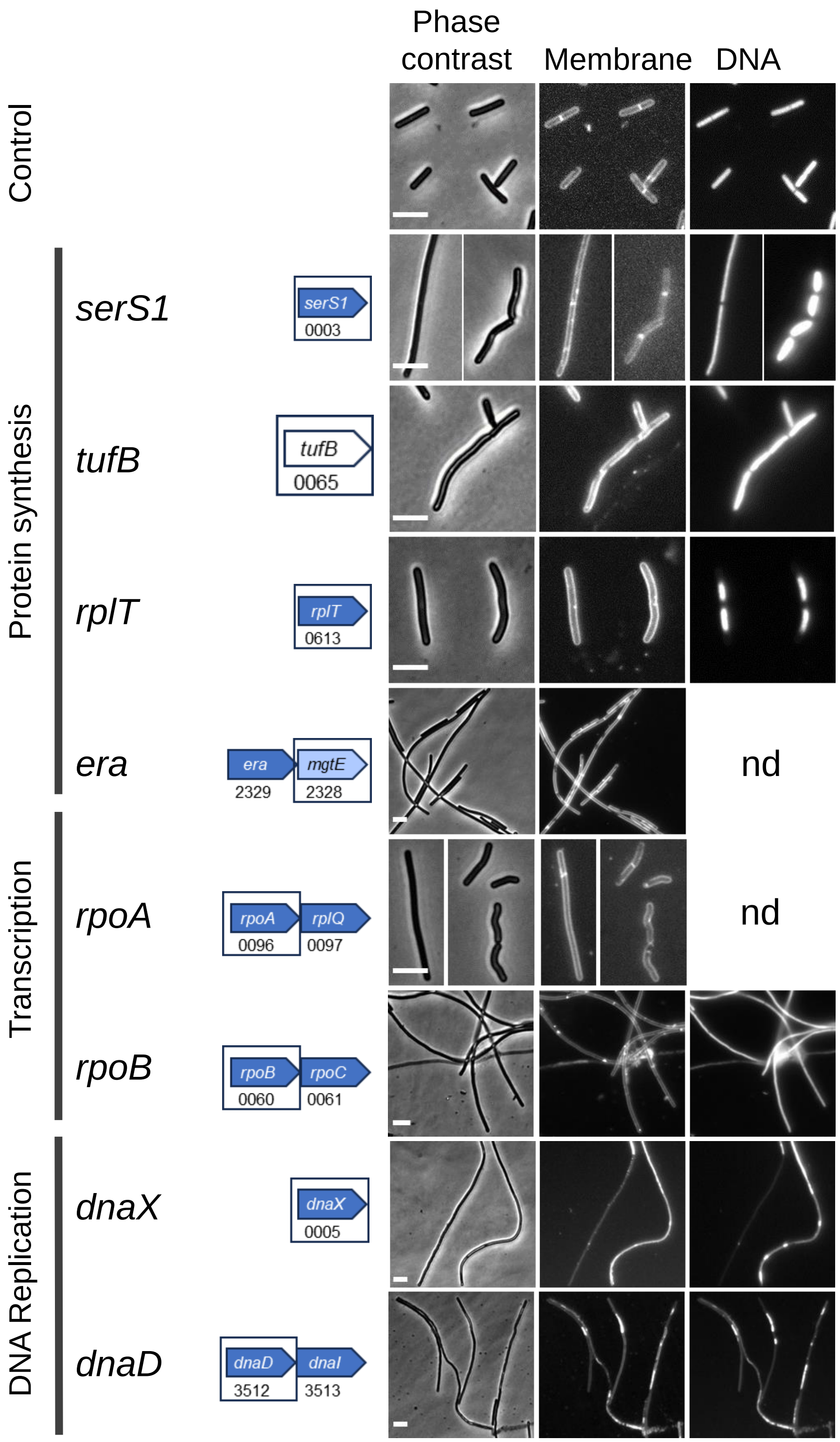
Phase contrast
Membrane
DNA
Control
Protein synthesis
serS1
serS1
0003
tufB
tufB
0065
rplT
rplT
0613
era
era
2329
mgtE
2328
nd
Transcription
rpoA
rpoA
0096
rplQ
0097
nd
rpoB
rpoB
0060
rpoC
0061
DNA Replication
dnaX
dnaX
0005
dnaD
dnaD
3512
dnaI
3513

Fig S1

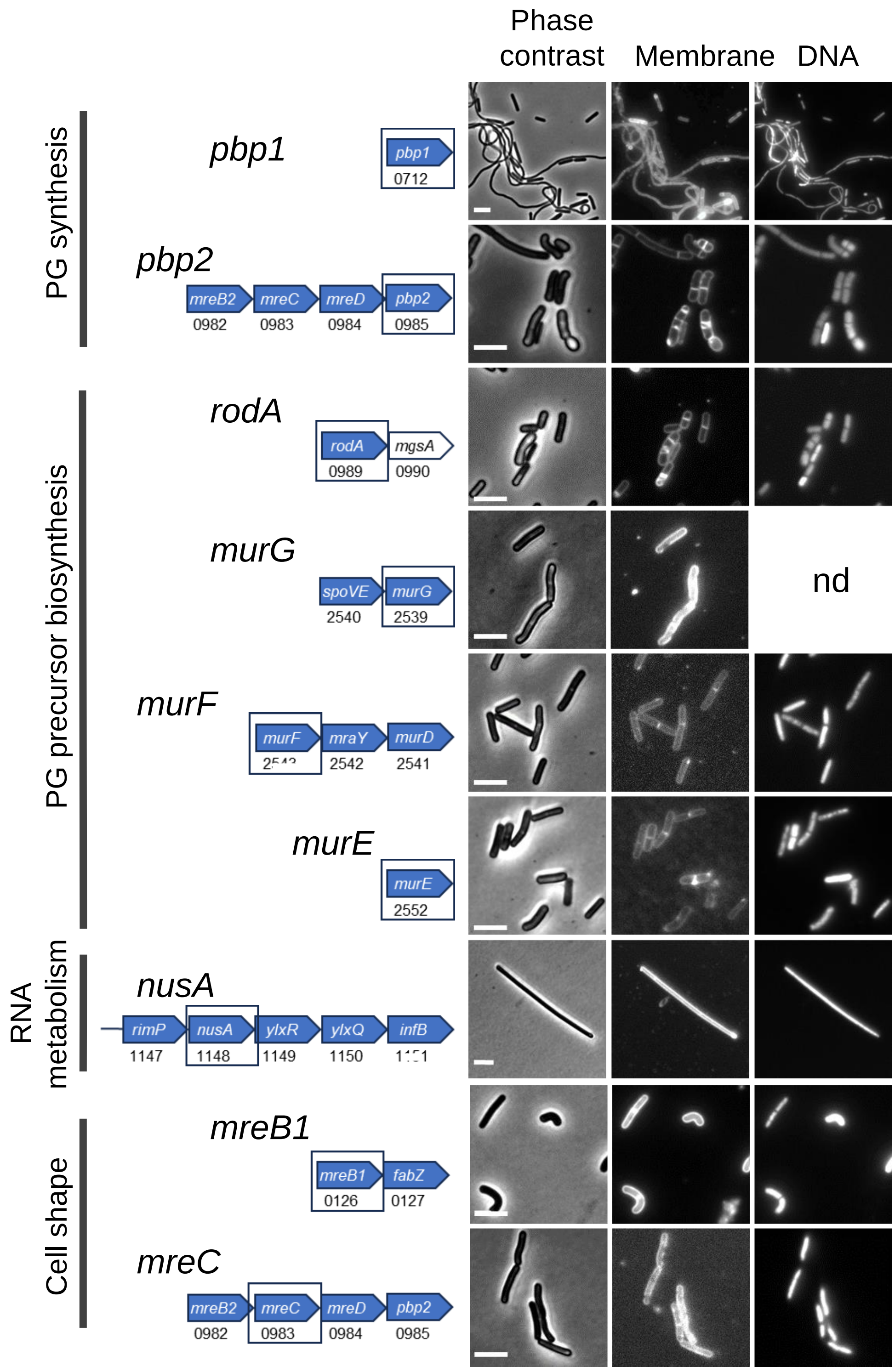
Phase contrast
Membrane
DNA
PG synthesis
pbp1
pbp1
0712
pbp2
mreB2
0982
mreC
0983
mreD
0984
pbp2
0985
PG precursor biosynthesis
rodA
rodA
0989
mgsA
0990
murG
spoVE
2540
murG
2539
nd
murF
murF
mraY
2542
murD
2541
murE
murE
2552
RNA metabolism
nusA
rimP
1147
nusA
1148
ylxR
1149
ylxQ
1150
infB
Cell shape
mreB1
mreB1
0126
fabZ
0127
mreC
mreB2
0982
mreC
0983
mreD
0984
pbp2
0985

Fig S1, cont.

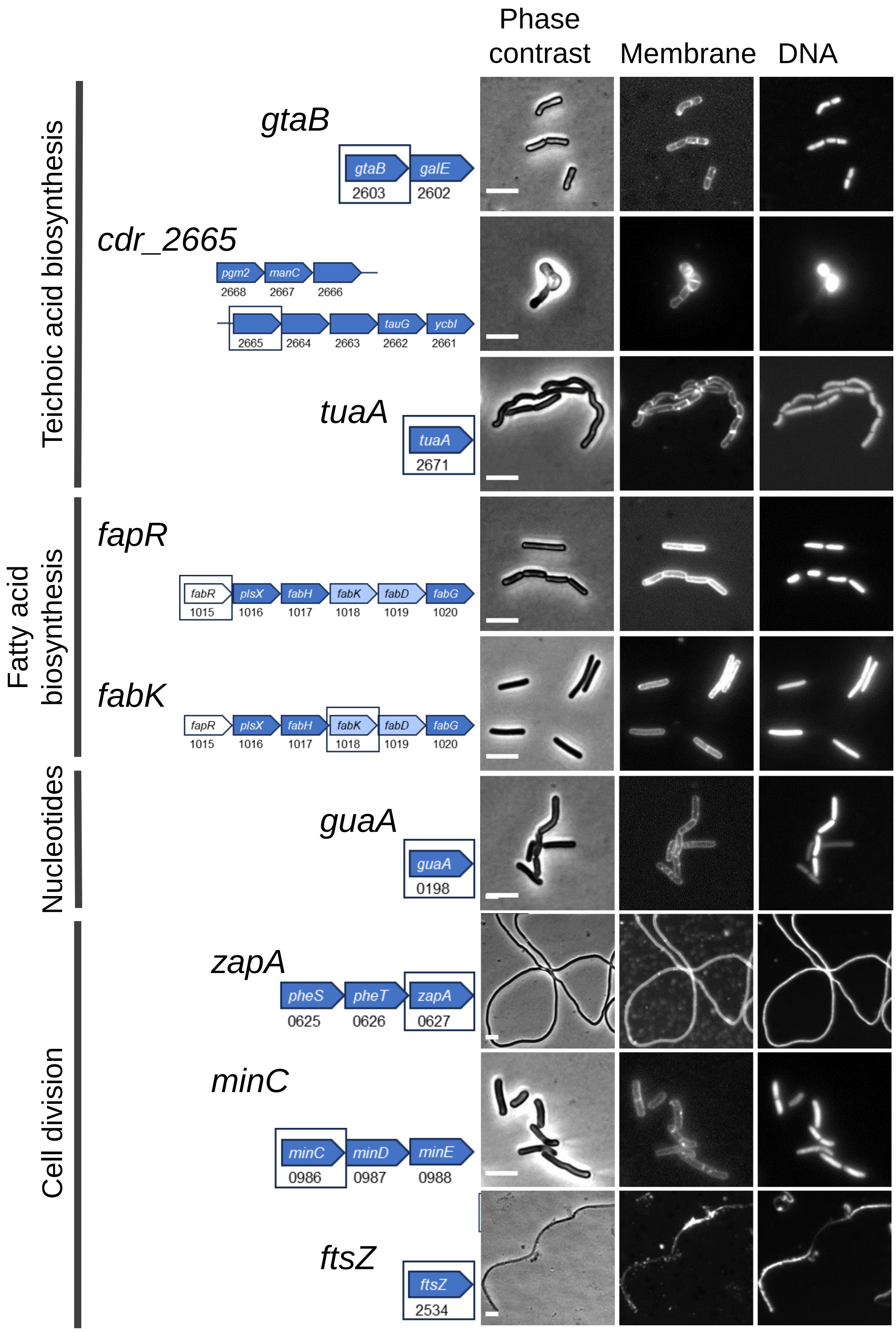

Phase contrast
Membrane
DNA
Teichoic acid biosynthesis
gtaB
gtaB
2603
galE
2602
cdr_2665
pgm2
2668
manC
2667
2666
2665
2664
2663
tauG
2662
ycbI
2661
tuaA
tuaA
2671
Fatty acid biosynthesis
fapR
fabR
1015
plsX
1016
fabH
1017
fabK
1018
fabD
1019
fabG
1020
fabK
fapR
1015
plsX
1016
fabH
1017
fabK
1018
fabD
1019
fabG
1020
Nucleotides
guaA
guaA
0198
Cell division
zapA
pheS
0625
pheT
0626
zapA
0627
minC
minC
0986
minD
0987
minE
0988
ftsZ
ftsZ
2534


Fig S1, cont.

**Figure S1. Extended version of Figure 2, morphology of CRISPRi strains with sgRNAs targeting genes in select functional pathways**. Left: pathway. Middle: Predicted transcription unit. Targeted genes are boxed and indicated above the operon diagrams. Numbers are R20291 locus tags. Genes are color-coded to indicate essentiality based on Tn-seq calls in Table S3. Blue: essential. Light blue: ambiguous. White: non-essential. Operon structure is not to scale. Right: Morphological changes based on phase contrast and fluorescence micrographs of cells scraped from viability plates. Membranes were stained with FM4-64 and DNA was stained with Hoechst 33342. n.d. means not determined. Size bars are 5 µm. Micrographs are representative of at least two experiments. The control strain expressed an sgRNA that does not target anywhere in the genome. Note: *tufB* is essential by CRISPRi but not by Tn-seq because the sgRNA against *tufB* also represses *tufA*.

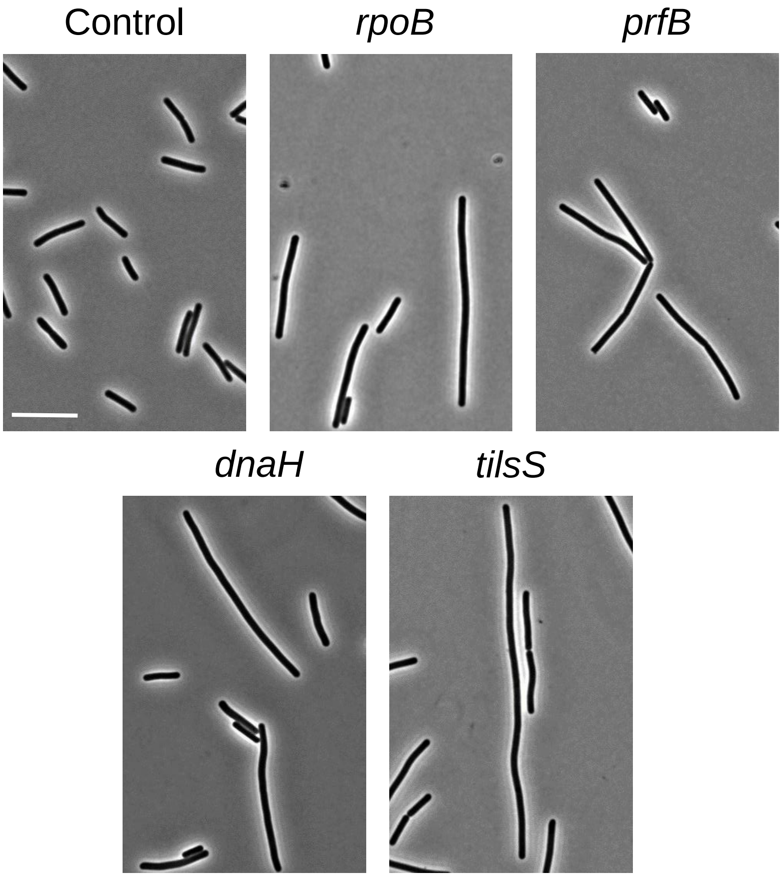


**Figure S2. Morphological phenotypes induced by CRISPRi against select genes in TY broth.** Starter cultures of the indicated CRISPRi strains that had been grown overnight in TY-Thi10 were diluted 1:400 into fresh TY-Thi10 with 1% xylose and grown for 5.5h (~ 6 mass doublings) to $OD_{600}$ ~0.35. Cells were fixed and photographed under phase contrast. Size bar = 5 µm. The control strain expressed an sgRNA against a sequence not present in R20291. Images are representative of at least two experiments.
