## Supplemental Tables 4-6 for "Analysis of Essential Genes in *Clostridioides difficile* by CRISPRi and Tn-seq"

Table S5: Plasmids

| Plasmid | Relevant features | Parent vector | Restriction enzymes to digest parent vector | PCR primers | PCR template | Assembly | Comments | Reference |
|---|---|---|---|---|---|---|---|---|
| pAP114 | *Pxyl::rfp* | | | | | | Genbank: MK368760 | Müh et al. 2019 |
| pCE1125 | *Pxyl::rfp-pbp1* | pDSW2040 | KpnI, SacI | 6660+6661 | pAP114 | HiFi[a] | | |
| pCE1126 | *Pxyl::rfp-pbp2* | pDSW2041 | KpnI, SacI | 6660+6661 | pAP114 | HiFi | | |
| pDSW1728 | *Ptet::rfp* | | | | | | Genbank: KT371995 | Ransom et al. 2015 |
| pDSW2021 | *Pxyl::* MCS-*rfp* | pAP114 | SacI, BamHI | P2382+P2383 | pDSW1728 | Ligation | | |
| pDSW2040 | *Ptet:rfp-pbp1* | pRAN473 | SphI, BamHI | P2401+P2402 | R20291 | Ligation | | |
| pDSW2041 | *Ptet:rfp-pbp2* | pRAN473 | SphI, BglII | P2303+P2304 | R20291 | Ligation | | |
| pHL07 | *Pxyl::rfp-cdr_2536(ylxX)* | pMK02 | KpnI, BamHI | 6687+6688 | R20291 | HiFi | | |
| pHL08 | *Pxyl::rfp-cdr_2537(ylxW)* | pMK02 | KpnI, BamHI | 6689+6690 | R20291 | HiFi | | |
| pHL09 | *Pxyl::rfp-cdr_2538(ftsQ)* | pMK02 | KpnI, BamHI | 6691+6692 | R20291 | HiFi | | |
| pHL13 | *Pxyl::rfp-cdr_3331* | pMK02 | KpnI, BamHI | 6699+6700 | R20291 | HiFi | | |
| pHL14 | *Pxyl::rfp-cdr_1285(ldt4)* | pMK02 | KpnI, BamHI | 6701+6702 | R20291 | HiFi | | |
| pHL15 | *Pxyl::rfp-cdr_2055(ldt5)* | pMK02 | KpnI, BamHI | 6703+6704 | R20291 | HiFi | | |
| pHL24 | *Pxyl::rfp-cdr_1165(ftsK)* | pMK02 | KpnI, BamHI | 6721+6722 | R20291 | HiFi | | |
| pIA131 | *Pxyl::* MCS-*rfp* | pDSW2021 | KpnI, BamHI | 6786+6788 | pMK02 | HiFi | Corrects mutation in pDSW2021 | |
| pIA33 | *$p_{xyl}$ R::dCas9-opt $P_{gdh}$::sgRNA-rfp catP* | | | | | | Parent plasmid for CRISPRi constructs; Genbank MK368761 | Müh et al. 2019 |
| pIA34 | *$p_{xyl}$ R::dCas9-opt $P_{gdh}$::sgRNA-neg catP* | | | | | | Negative control CRISPRi plasmid | Müh et al. 2019 |
| pJR01 | *Pxyl::cdr_2503(divIVA)-rfp* | pIA131 | KpnI, SacI | 6675+6676 | R20291 | HiFi | | |
| pJR05 | *Pxyl::rfp-cdr_2505(ylmG)* | pMK02 | KpnI, BamHI | 6683+6684 | R20291 | HiFi | | |
| pJR06 | *Pxyl::cdr_2506(sepF)-rfp* | pIA131 | KpnI, SacI | 6685+6686 | R20291 | HiFi | | |
| pJR17 | *Pxyl::rfp-cdr_0983(mreC)* | pMK02 | KpnI, BamHI | 6707+6708 | R20291 | HiFi | | |
| pJR18 | *Pxyl::rfp-cdr_0984(mreD)* | pMK02 | KpnI, BamHI | 6709+6710 | R20291 | HiFi | | |
| pJR23 | *Pxyl::rfp-cdr_2283(mgt)* | pMK02 | KpnI, BamHI | 6731+6732 | R20291 | HiFi | | |
| pLD1 | *Pxyl::rfp-cdr_2797(ldt1)* | pMK02 | KpnI, BamHI | 6938+6939 | R20291 | HiFi | | |
| pLD7 | *Pxyl::rfp-cdr_2534(ftsZ)* | pMK02 | KpnI, BamHI | 6998+6999 | R20291 | HiFi | | |
| pMK02 | *Pxyl::rfp-cdr_0018* | pCE1125 | SphI-BamHI | 6578+6579 | R20291 | HiFi | | |
| pMK50 | *Pxyl::rfp-cdr_0989(rodA)* | pMK02 | KpnI, BamHI | 7672+7673 | R20291 | HiFi | | |
| pRAN473 | *Ptet::rfp-* MCS | | | | | | | Ransom et al. 2015 |
| pRPF215 | Ptet-himar1-Ter(slpA)-transposon | | | | | | | Dembek et al. 2015 |

[a]: HiFi assembly (NEB)

Table S4: Strains

| Strain | Genotype | Comment | Source |
|---|---|---|---|
| *E. coli* | | | |
| HB101/pRK24 | F− *mcrB mrr hsdS20* ($r_B^-$ $m_B^-$) *recA13 leuB6 ara-14 proA2 lacY1 galK2 xyl-5 mtl-1 rpsL20* | | Dineen et al. 2007, Trieu-Cuot et al. 1991 |
| *C. difficile* | | | |
| R20291 | Wild-type *C. difficile* strain from UK outbreak (ribotype 027) | | |
| UM1320 | R20291/pCE1125 | *Pxyl::rfp-pbp1 catP* | This study |
| UM1175 | R20291/pCE1126 | *Pxyl::rfp-pbp2 catP* | This study |
| KB412 | R20291/pHL07 | *Pxyl::rfp-ylxX catP* | This study |
| KB413 | R20291/pHL08 | *Pxyl::rfp-ylxW catP* | This study |
| KB418 | R20291/pHL09 | *Pxyl::rfp-ftsQ catP* | This study |
| KB448 | R20291/pHL13 | *Pxyl::rfp-cdr_3331 catP* | This study |
| KB449 | R20291/pHL14 | *Pxyl::rfp-ldt4 catP* | This study |
| KB455 | R20291/pHL15 | *Pxyl::rfp-ldt5 catP* | This study |
| KB454 | R20291/pHL24 | *Pxyl::rfp-ftsK catP* | This study |
| UM0359 | R20291/pIA34 | *pxylR::dCas9-opt Pgdh::sgRNA-neg catP* | Müh et al. 2019 |
| UM1247 | R20291/pJR01 | *Pxyl::divIVA-rfp catP* | This study |
| UM1251 | R20291/pJR05 | *Pxyl::rfp-ylmG catP* | This study |
| UM1252 | R20291/pJR06 | *Pxyl::sepF-rfp catP* | This study |
| UM1254 | R20291/pJR17 | *Pxyl::rfp-mreC catP* | This study |
| UM1255 | R20291/pJR18 | *Pxyl::rfp-mreD catP* | This study |
| UM1258 | R20291/pJR23 | *Pxyl::rfp-mgt catP* | This study |
| LD0110 | R20291/pLD01 | *Pxyl::rfp-ldt1 catP* | This study |
| LD0122 | R20291/pLD07 | *Pxyl::rfp-ftsZ catP* | This study |
| UM1420 | R20291/pMK50 | *Pxyl::rfp-mrdB catP* | This study |
| CDE2716 | R20291/pRPF215 | *Ptet-himar1-Ter(slpA)-transposon* | Dembek et al. 2015 |

Table S6: Oligonucleotides

| Oligo | Sequence | Comment |
|---|---|---|
| CDEP6578 | TATGGATGAATTATATAAAATGAACGGTGGTGGTGGTGGTACCTTGATAAGGAAAGTAACAAGAATATTG | |
| CDEP6579 | ATTTAAAGTTTTATTAAAACTTATAGGATCCTTAATTTTTAGGTTTTGCAAATATCATTC | |
| CDEP6660 | GCTTCTTATTTTTATGGTACCGTATAATTTAAAGGGTTATGGCTGT | |
| CDEP6661 | TCTCCTTTACTGCAGGAGCTCACTTCATAACTATCGTTTTCCT | |
| CDEP6675 | CGATAGTTATGAAGTGAGCTCAAGGAGAAAATTTTATGCTAACTCCAATTGAGATAGAAAATAAAG | |
| CDEP6676 | AGATACCATAGATCCGGTACCAGATCCAGATCCTTCTAAAGTTGTAGCAGCTTCATCATTACTATA | |
| CDEP6683 | AACGGTGGTGGTGGTGGTACCATGGGAACAATAAGAATTGCAC | |
| CDEP6684 | ATTTAAAGTTTTATTAAAACTTATAGGATCCTTAGAATATAACCAATACTAATTGTCTTAGG | |
| CDEP6685 | CGATAGTTATGAAGTGAGCTCAAGGAGAAAATTTTATGTCAAATGGAATAATATCTAAATTTAAGAATTG | |
| CDEP6686 | AGATACCATAGATCCGGTACCAGATCCAGATCCTTTATTTTGCCAAGGGAAGAATG | |
| CDEP6687 | AACGGTGGTGGTGGTGGTACCATGAAAAAGAAAATAGTTATATTAGGAGCA | |
| CDEP6688 | ATTTAAAGTTTTATTAAAACTTATAGGATCCTTATTTTGCAAATAAATTCACTTTATTCCC | |
| CDEP6689 | AACGGTGGTGGTGGTGGTACCTTGAAAAAAAGATTTATAAATAAAAGTATAATCATAGG | |
| CDEP6690 | ATTTAAAGTTTTATTAAAACTTATAGGATCCTTATTTCCCCTCTGCTTTGTTTATATC | |
| CDEP6691 | AACGGTGGTGGTGGTGGTACCTTGAAAAAAAGAAGAAAGTTAAATACGAATAATG | |
| CDEP6692 | ATTTAAAGTTTTATTAAAACTTATAGGATCCCTATCCTTCTGGACTATACACTGC | |
| CDEP6699 | AACGGTGGTGGTGGTGGTACCATGGAACAGAACATAACTTTGAAAG | |
| CDEP6700 | ATTTAAAGTTTTATTAAAACTTATAGGATCCCTATCTAAATACCTTTTTCAAGAAACTC | |
| CDEP6701 | AACGGTGGTGGTGGTGGTACCATGAGTAATGTGAACAAAAAGCTAG | |
| CDEP6702 | ATTTAAAGTTTTATTAAAACTTATAGGATCCTTAAGCTTGCGGTTGTGG | |
| CDEP6703 | AACGGTGGTGGTGGTGGTACCATGGGCAGAAGAACAACAAG | |
| CDEP6704 | ATTTAAAGTTTTATTAAAACTTATAGGATCCCTATTTTTTTAATTCTTTATAAGAACTATTTGATATATG | |
| CDEP6707 | AACGGTGGTGGTGGTGGTACCTTGGTGATGGCCTTGAGATTTG | |
| CDEP6708 | ATTTAAAGTTTTATTAAAACTTATAGGATCCTTATCCTATATTTCTTGGTTCTATGACTA | |
| CDEP6709 | AACGGTGGTGGTGGTGGTACCATGAAAAAAGTTTTACTTTGTCTGTTG | |
| CDEP6710 | ATTTAAAGTTTTATTAAAACTTATAGGATCCCTAATCTTCTTTAAGTTTTAAAACTGCTC | |
| CDEP6721 | AACGGTGGTGGTGGTGGTACCATGGCCACGAAAAAGAAGAA | |
| CDEP6722 | ATTTAAAGTTTTATTAAAACTTATAGGATCCCTATTCACCTTCTAAATTCTGCAAATC | |
| CDEP6731 | AACGGTGGTGGTGGTGGTACCATGGGTTATTACAATGATAATGATAATAATAAAATAAG | |
| CDEP6732 | ATTTAAAGTTTTATTAAAACTTATAGGATCCTTAATCAATTTCTGATTTAAATGGAGTG | |
| CDEP6786 | CTCTCTAGAGTCGACGGTACCGGATCTATGGTATCTAAAGGAGAAGAAGATAATATG | |
| CDEP6788 | CTTTTTCTATTTAAAGTTTTATTAAAACTTATAGGATCCTTATTTATATAATTCATCCATACCTCCTGTTG | |
| CDEP6938 | AACGGTGGTGGTGGTGGTACCATGATTGATGGAAAAGAGGTAAAAC | |
| CDEP6939 | ATTTAAAGTTTTATTAAAACTTATAGGATCCTAGTATAAAATAATTGGTGTACCTGG | |
| CDEP6998 | AACGGTGGTGGTGGTGGTACCATGATGCTAAACTTTGACGTAGA | |
| CDEP6999 | ATTTAAAGTTTTATTAAAACTTATAGGATCCTTATCTTCTTCTTCTTAGGAATGTAGG | |
| CDEP7672 | AACGGTGGTGGTGGTGGTACCTTGAAAAAAATCAATATTAATTATAAGTCGAC | |
| CDEP7673 | TTTAAAGTTTTATTAAAACTTATAGGATCTTAGAAATTTATTTTCTTTCTTCTCATACAC | |
| P2303 | CACGCATGCGGTGGATCTGAAGAAAATAAACAAGTTGATAGACTAAAAATAATTAAG | |
| P2304 | CACAGATCTAAGCTATCTATTAATTTGTGACTCG | |
| P2401 | CAC GCATGCGGTGGATCTAATAACGATAATGATAAAAATGGAAATAAAATCAGG | |
| P2402 | CACGGATCCTTAAGTTGCTGGAGTTCCACC | |
| P5 Tn Px | AATGATACGGCGACCACCGAGATCTACACTCTTTCCCTACACGACGCTCTTCCGATCTNNNNAGATCA [illegible] | **NNNN**: variable barcode |
| P7 16G | CAAGCAGAAGACGGCATACGAGCTCTTCCGATCTGGGGGGGGGGGGGGGG | |
| Tn-ermB-2 | ATCACTCCTTCTTAATTACAAATTTTTAGCATCTAATTTAACTTCAATTCCTATTATAC | |

**References Supplemental Information**

1. Dembek M, Barquist L, Boinett CJ, Cain AK, Mayho M, Lawley TD, Fairweather NF, Fagan RP. 2015. High-throughput analysis of gene essentiality and sporulation in *Clostridium difficile*. MBio 6:e02383.
2. Dineen SS, Villapakkam AC, Nordman JT, Sonenshein AL. 2007. Repression of *Clostridium difficile* toxin gene expression by CodY. Mol Microbiol 66:206-19.
3. Edwards AN, Wetzel D, DiCandia MA, McBride SM. 2022. Three Orphan Histidine Kinases Inhibit *Clostridioides difficile* Sporulation. J Bacteriol 204:e0010622.
4. Elfmann C, Dumann V, van den Berg T, Stülke J. 2025. A new framework for SubtiWiki, the database for the model organism *Bacillus subtilis*. Nucleic Acids Res 53:D864-d870.
5. Koo BM, Kritikos G, Farelli JD, Todor H, Tong K, Kimsey H, Wapinski I, Galardini M, Cabal A, Peters JM, Hachmann AB, Rudner DZ, Allen KN, Typas A, Gross CA. 2017. Construction and Analysis of Two Genome-Scale Deletion Libraries for *Bacillus subtilis*. Cell Syst 4:291-305.e7.
6. Müh U, Pannullo AG, Weiss DS, Ellermeier CD. 2019. A Xylose-Inducible Expression System and a CRISPR Interference Plasmid for Targeted Knockdown of Gene Expression in *Clostridioides difficile*. J Bacteriol 201:e00711-18.
7. Ransom EM, Ellermeier CD, Weiss DS. 2015. Use of mCherry Red fluorescent protein for studies of protein localization and gene expression in *Clostridium difficile*. Appl Environ Microbiol 81:1652-60.
8. Shrestha S, Taib N, Gribaldo S, Shen A. 2023. Diversification of division mechanisms in endospore-forming bacteria revealed by analyses of peptidoglycan synthesis in *Clostridioides difficile*. Nat Commun 14:7975.
9. Trieu-Cuot P, Carlier C, Poyart-Salmeron C, Courvalin P. 1991. Shuttle vectors containing a multiple cloning site and a lacZ alpha gene for conjugal transfer of DNA from *Escherichia coli* to gram-positive bacteria. Gene 102:99-104.
10. Underwood S, Guan S, Vijayasubhash V, Baines SD, Graham L, Lewis RJ, Wilcox MH, Stephenson K. 2009. Characterization of the sporulation initiation pathway of *Clostridium difficile* and its role in toxin production. J Bacteriol 191:7296-305.
11. van Eijk E, Paschalis V, Green M, Friggen AH, Larson MA, Spriggs K, Briggs GS, Soultanas P, Smits WK. 2016. Primase is required for helicase activity and helicase alters the specificity of primase in the enteropathogen *Clostridium difficile*. Open Biol 6.
